# Sampling Protein Language Models for Functional Protein Design

**DOI:** 10.1101/2025.09.14.676087

**Authors:** Jeremie Theddy Darmawan, Yarin Gal, Pascal Notin

## Abstract

Protein language models have emerged as powerful tools for learning rich protein representations, improving performance in tasks like structure prediction, mutation effect estimation, and homology detection. Their ability to model complex sequence distributions also holds promise for designing novel, functional proteins with broad applications in therapeutics, materials, and sustainability. However, due to the vastness of sequence space, efficient exploration methods are essential for protein engineering. Despite this, most existing protein design approaches using protein language models rely on single-mutant sampling strategies borrowed from Natural Language Processing, which fail to capture critical epistatic interactions between amino acid positions that are essential for protein function. In this work, we develop a comprehensive in silico protein design evaluation framework to systematically compare different sampling methods. After a thorough review of existing sampling strategies for language models, we introduce several approaches specifically tailored for protein design. We demonstrate that sampling strategies that consider multiple mutations simultaneously significantly outperform single-mutant approaches by better capturing epistatic effects between residue pairs. We evaluated these strategies using our framework, investigating the effects of key hyperparameters and providing practical guidance on the relative strengths of each method depending on design objectives.

## 1 INTRODUCTION

Proteins are complex macromolecules that perform a wide diversity of essential functions in biological systems, such as catalyzing biochemical reactions, facilitating cellular signaling, and providing structural support. They cover a vast sequence space, representing all possible variations of amino acid chains that can fold into functional structures. Despite significant advancements in sequencing technologies, the protein sequences currently known and characterized represent only a small fraction of this immense sequence space^1^. Protein design methodologies seek to efficiently explore this vast space to create novel protein sequences. This exploration can be achieved either by iteratively mutating existing functional sequences of interest^2–4^ or by designing entirely new sequences from scratch, drawing on existing templates and biochemical constraints^5,6^. Protein language models (pLMs) have emerged as powerful tools to learn complex distributions over protein sequences, leading to superior performance on a broad range of downstream tasks, such as structure prediction^7,8^, fitness and mutation effects prediction^9–12^, homology detection^13^, or viral evolution^14,15^. As generative models of protein sequences, pLMs are ideally suited to sample novel functional proteins that have not yet been observed, making them invaluable in supporting protein engineering workflows. Given the massive size of the corresponding sample space, together with the diversity of protein design objectives, the success of pLM-based protein engineering efforts depends not only on ever-improving models but also on efficient methods to *sample* from these models. While the field has witnessed the emergence of several pLMs for protein design^16–20^, efficient sampling methods have been understudied to date. Prior works have so far primarily leveraged sampling strategies initially developed for Natural Language Processing (NLP) tasks (§ 2.1), with limited insight into the impact of various strategies on the properties of the generated sequences, and without taking advantage of the unique characteristics of protein sequences relative to natural language sequences.

In this work, we sought to develop and explore different methods for sampling from pLMs, systematically analyzing the characteristics of the generated sequences based on the chosen sampling strategy or core underlying hyperparameters. To that end, we first devise a robust *in silico* evaluation framework to benchmark different protein design methods. While our focus is on pLMs in general, we place a greater emphasis on autoregressive pLMs, given their ability to support a broader range of sampling methods (§ 3.1). We also place ourselves in the zero-shot design setting, aiming to design novel functional sequences without relying on experimental labels acquisition. Our contributions are as follows. We introduce a high-level framework and taxonomy to describe the landscape of existing pLM sampling methods (§ 3.2). We develop a suite of efficient sampling strategies tailored to protein design objectives: High-Probability Filter (HPF), Quantitative-Function Filter (QFF), Attention-Matrix Sampling (AMS), Random Stratified Filtering (RSF), and Monte Carlo Tree Search (MCTS) schemes (§ 3.3). We devise a holistic framework for the *in silico* evaluation of protein engineering methods, spanning core objectives such as functional relevance, sequence diversity, and fitness (§ 3.4). We conduct an in-depth comparison of the various pLM sampling methods, analyze the effect of key sampling hyperparameters on performance, and discuss the relative strengths of the different methods depending on design objectives (§ 4). We open-source our codebase to provide convenient access to our *in silico* generation and evaluation framework, including all sampling methods, in a unified interface: https://github.com/i3LBI19-OATML/sampling_plm.

## 2 BACKGROUND AND RELATED WORKS

### 2.1 Sampling Language Models

Language models are generative models seeking to approximate a distribution over sequential inputs. Once trained, we can sample from these generative models to create new sequences that may not have been part of the initial training data. In NLP, since the creation of novel sequences may be driven by various objectives (e.g., creativity, coherence, fluency, factual information), a wide range of sampling strategies has been developed to promote certain characteristics of generated sequences^21^. Throughout the text, we refer to these sampling approaches as the *standard sampling strategies*. We describe below some of these strategies, where *x* represents the sequence token. The simplest strategy is random sampling, which samples items based on their probabilities as provided in Equation (1), where 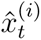 is the unnormalized logits at step *t* for item *i* in the vocabulary *V*, divided by the smoothing temperature parameter *T* which controls the flatness of the distribution.

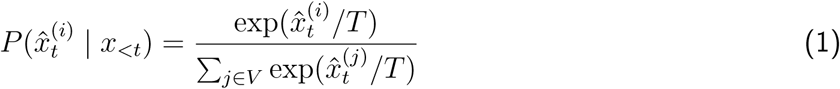

In order to avoid sampling items with extremely low probability under the language model, Top-k sampling selects the subset *V*^(*k*)^ of the *k* elements with highest probabilities, sets the probabilities of the other items to zero, and renormalizes probabilities for the Top-k items as expressed in Equation (2).

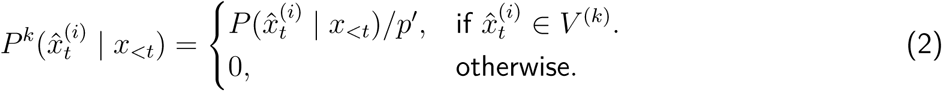

where *x_<t_* corresponds to sequence items previously generated, and 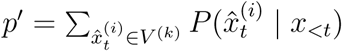 is the probability re-normalization factor. **Greedy sampling** is an extreme version of Top-k sampling that only takes the item with the highest (top-1) probability in the distribution.

In practice, using a fixed-size k may be suboptimal when the flatness of the distribution varies significantly across generated tokens in the sequence: in some instances, relevant items will be censored, while in others certain low-likelihood items may be sampled. To address this problem, Top-p sampling ^22^ (or nucleus sampling) first computes the corresponding cumulative distribution and censors it as soon as the CDF exceeds *p*, giving rise to the set *V^(p)^* of all non-censored items. Mathematically, this leads to a censored probability analogous to the one in Equation (2), except for the renormalization factor 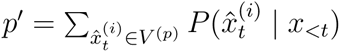.

**Beam search** views a sequential sampling process as a search problem exploring the most promising mutations at each step in a tree-like structure, maintaining a fixed number of possible solutions called “beam width”. Equation (3) outlines this approach, where elements *x_t_* of the beam width at step *t* are the ones that maximizes the scoring function *s*(.) (typically the likelihood) of the concatenation (denoted by the operator [:]) of sequences *x_<t_* in the beam width *X_<t_* at the prior step, together with any other element 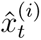 from the vocabulary *V* at step t.

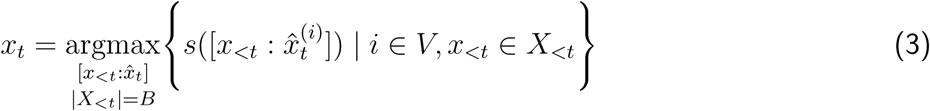

Other sampling techniques maximizing human expectations and preferences were also developed for NLP. Typical sampling attempts to balance out the information content of the generated sequence and the expected information content (entropy) as shown in Equation (4)^23^. The truncation set *C*(*x_<t_*) represents the solution to the minimization problem, P is the power set operator, *V* is the vocabulary, *H* is the entropy, and *τ* is the amount of probability mass to be considered.

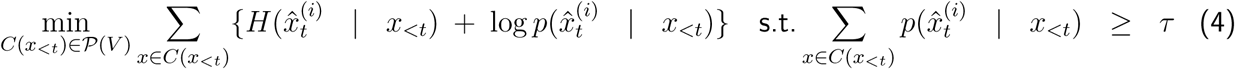

Basu et al. ^24^ developed a fine-tuning-free method, called mirostat sampling, which self-adapts the perplexity of generated sequence at a predefined level based on human preference. This is done by truncating the token candidate list using top-*k*, while self-adaptation can be achieved by adjusting the error based on the observed and target perplexity difference. Algorithm S1 details this approach, where S(*x*) is the observed perplexity, *τ* is the target perplexity, *m* is the number of most probable tokens, *µ* is the maximum perplexity, *η* is the learning rate, *ɛ* is the error term, and *s*^ refers to Zipf’s exponent^24^.

Language modeling can also be viewed as a sequence of decisions where the current generated tokens (state) will influence the next tokens (moves) being generated to achieve certain objectives. Monte Carlo Tree Search (MCTS)^25,26^ has been used in machine translation and, as a result, for protein design^27,28^. Recently, Wang et al. ^28^ leveraged a MCTS-based approach to carry out a 35 residue-restricted iterative protein design. The search mechanism is based on iterative rounds of simulations from the current state where the algorithm will play out subsequent moves. It leverages a UCT (Upper Confidence bounds applied to Trees) algorithm to decide between exploring or exploiting the current state. Exploration refers to playing moves on unexplored nodes whereas exploitation re-investigates nodes that have been previously played for higher rewards. The information on each node visits (potential mutations) and rewards (improvement in fitness) is backpropagated through the tree to improve subsequent simulation attempts. Equation (5) expresses the UCT algorithm for a particular node, where *W_i_* is the total number of successful (positive) simulations, *S_i_* is the total number of simulations, *S_p_* is the total number of simulations of the parent node, and *c* is the exploration parameter.

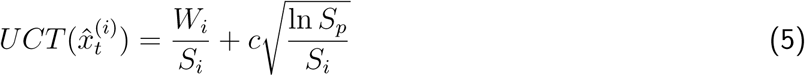

### 2.2 Protein Language Models

The majority of protein language models can be classified into two broad categories: autoregressive (AR) and masked-language models (MLM), although other learning paradigms such as seq2seq^29^ or FIM^13^ have also been explored. Unirep^16^ was the first pLM trained autoregressively across unaligned protein sequences across protein families. With the improvements seen in Transformers^30^ for NLP, masked-language models such as ProteinBERT, ESM-1b, TAPE-BERT, MSA Transformer, and SaProt^10,12,31–33^, as well as autoregressive models, such as ProtGPT2, Progen, Tranception, RITA, ZymCTRL, PoET^17,18,20,34–36^, have been proposed to further increase the quality of learned representations. These pLMs have been shown to achieve remarkable performance across diverse tasks, such as fitness prediction^9,34^ or structure prediction^7^. For protein design, prior pLM works have mainly relied on autoregressive generation with simple sampling methods discussed in § 2.1, such as Top-k, Top-p, random, or greedy sampling ^17,19,20,37,38^. Sgarbossa et al. ^38^ developed a method that iteratively applies random masking of residues in the protein sequence and fills them greedily based on MSA Transformer^33^ predictions. Diffusion models have also been applied for protein design through this mechanism^39–41^. While these various approaches were successful in generating novel protein sequences, none of these prior works have systematically studied the effects and relative trade-offs of various sampling methods on the properties of generated sequences.

## 3 METHODS

### 3.1 Taxonomy for pLM sampling methods

Sampling strategies for ML-based protein design can be broadly grouped into either Iterative Redesign Sampling (IRS) or AutoRegressive Sampling (ARS), as depicted in Figure 1. We can leverage Protein Language Models (pLMs) to generate novel protein sequences using either of these approaches^42^. Additionally, any of the sampling methods outlined in § 2.1 can be applied in conjunction with both IRS and ARS approaches.

**Figure 1:**
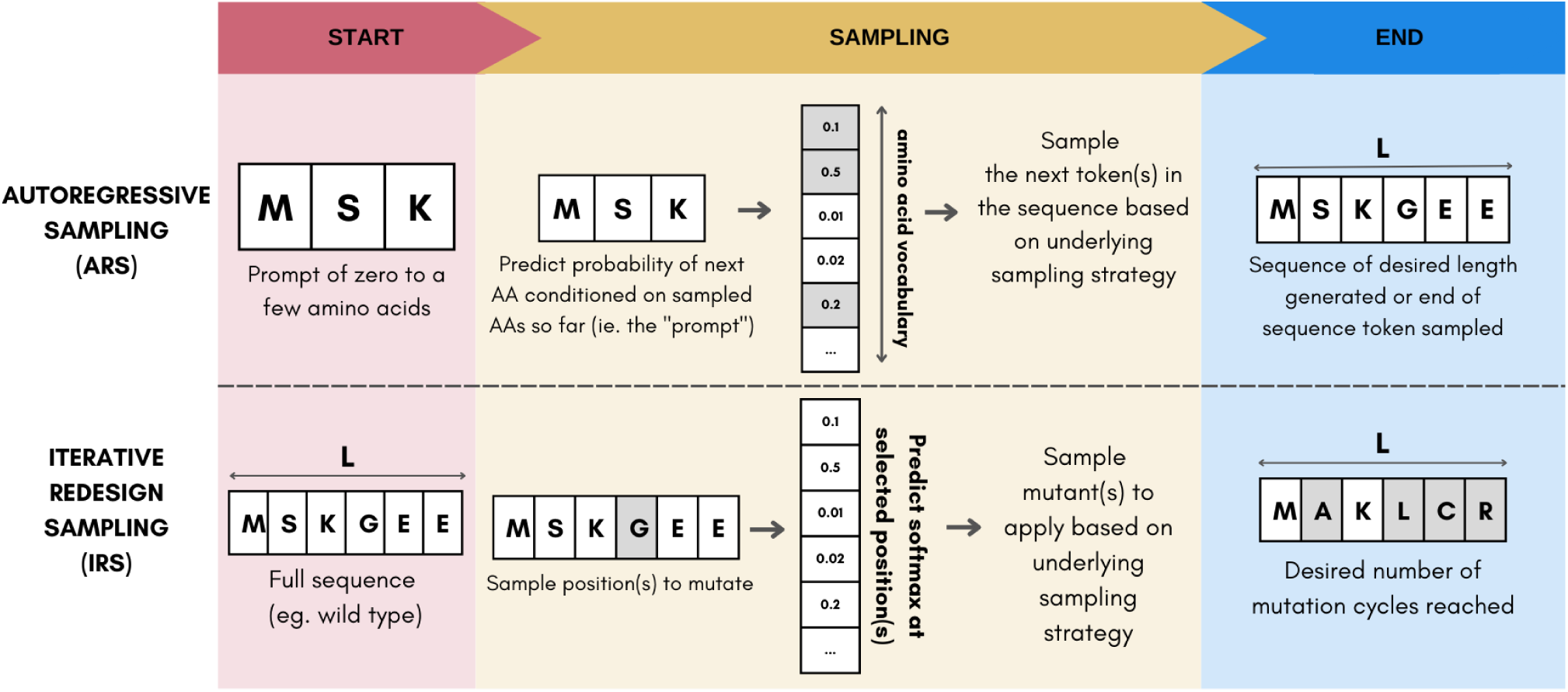
Taxonomy of Protein Language Models sampling methods. We discriminate between two strategies for sampling pLMs: Autoregressive Sampling and Iterative Redesign Sampling.

#### AutoRegressive Sampling

Given an initial residue prompt, ARS sampling methods iteratively extend the amino acid (AA) sequence by one or multiple residues at each step. ARS strategies can be conditioned with a prompt containing a few residues, or be fully unconditional (empty prompt). A prompted generation typically allows for more targeted and family-specific generation^19^. The process stops when the desired sequence length has been reached or an ‘end of sequence’ token is generated. Pseudocode for the approach is provided in Algorithm S8.

#### Iterative Redesign Sampling

IRS-based sampling starts by taking a template protein sequence (e.g., a natural protein sequence) and generating all possible single (or a subset of multiples) mutations, either at every position or at specific Position(s) of Interest (PoI). At each iteration, one (or multiple) substitution mutations are introduced and the process stops once a predefined number of evolution cycles is achieved. IRS encompasses Markov Chain Monte Carlo (MCMC) approaches such as Gibbs sampling^38,43–46^. Each optimal mutation is determined by a fitness score derived from the underlying pLM and a predefined sampling strategy. This strategy can be applied not only to traditional MLM pLMs, but also to newer masked diffusion language modeling pLMs. An illustration of the code for single-mutant IRS is provided in Figure S1 and the detailed algorithm is available in Algorithm S7.

### 3.2 Single-Mutant Protein Generation

Methods in this section focus on sampling a single mutant at each iteration. Single mutant IRS (IRS-singles) strategies require a template sequence as the initial starting point. They subsequently generate all possible single amino acid substitutions at each position, which are then scored with a fitness predictor to select which of these possible mutations to apply. A predefined number of iterative cycles specifies when to stop the entire process. Conversely, single mutant ARS (ARS-singles) strategies iteratively build up a full sequence by adding a single residue at each cycle. This residue is sampled autoregressively out of all 20 possible amino acids with the underlying generative model. In both ARS and IRS single strategies, we sample with one of the standard approaches introduced in § 2.1, including random, greedy, Top-k, Top-p, typical, and mirostat sampling. Single-mutant-based methods are straightforward to instantiate and have a linear computational complexity in the sequence length.

### 3.3 Multiple-Mutant Protein Generation

While simpler to implement and computationally tractable, methods that sample a single mutant at each iteration may result in suboptimal design performance as they ignore potentially critical epistatic effects between positions^43,47^. Uncovering these relationships requires to consider the interactions between multiple mutated positions simultaneously^48^. However, given the combinatorial number of such interactions, an exhaustive enumeration and scoring of all possible mutants is impractical. Efficient multi-mutant strategies thus rely on simplifying assumptions to make that search tractable. In this work, we introduce several such strategies that focus on efficient sampling of pairs of mutants. By adjusting the key variable *N* in the following strategies, we control the number of proteins explored at each step. In all our experiments, we set *N* = 96, as previously identified to be the minimal for exploration^37^. We refer to these methods as ARS-doubles and IRS-doubles. We provide a high-level illustration for the IRS-doubles strategies in Figure S1 and detail the corresponding pseudocode in Algorithm S9.

#### Quantitative-Function Filter (QFF)

We first generate all possible mutant pairs. We then quantify the protein fitness for all possible pairs with a Potts model, EVmutation^49^. This significantly reduces the computational complexity compared with doing the same with a large pLM, while keeping the ability to quantify epistatic effects between residue pairs. We then select the top-N predicted mutations, score these with the pLM. Lastly, we sample the double mutant to apply out of these N options with one of the sampling strategies discussed in § 2.1.

#### High-Probability Filter (HPF)

We first generate and score all possible single amino acid substitutions. The Top-*k* (e.g., *k* = 100) single mutants with the highest fitness scores are then selected and combined together to generate a set of high-quality double mutants. This drastically reduces the number of double mutations to consider, reducing the overall computational complexity of the algorithm to being linear in sequence length. From this set, we randomly sample *N* mutant pairs which we score with the pLM. While it is possible to focus on the mutant pairs with the highest average scores, we instead randomly sample a number N of them to promote exploration in sequence space. The double mutant to apply in the current cycle is then sampled out of these *N* options with one of the standard sampling strategies discussed in § 2.1.

#### Attention-Matrix Sampling (AMS)

We first score all possible single mutants and select the top-*K* with the highest predicted fitness. Using the same pLM, we identify PoIs as residues with the highest average attention values in the self-attention layers. From this PoI set, we generate all possible mutant pairs and score them using a lightweight fitness model (EVmutation), selecting the top-*N* for further evaluation with the pLM. A double mutant is then sampled from these *N* options using a standard strategy. Alternative selection methods, such as identifying spatially proximal residues from experimental or predicted protein structures, may also be used instead of attention scores.

#### Random-Stratified Filter (RSF)

All possible double mutations are generated from the top-*K* single mutants. Each sequence is scored with a lightweight fitness prediction model (EVmutation) and stratified into four bins based on their scores: {*S*_very low_*, S*_low_*, S*_high_*, S*_very high_}. We then take a total of *N* double-mutants by taking the top-*N/*4 samples from each stratum to increase the diversity of mutations, intuitively mutants in different EVmutation score bins will be non-redundant and diverse in sequence space. This introduces mutation diversity at an intermediate level before further protein scoring. Lastly, we sample the double mutant to apply out of these N options with one of the standard sampling strategies.

#### Monte Carlo Tree Search (MCTS)

We start from a full sequence (natural or mutated) and consider each of its residues as a potential parent node^27,28^. In each search round, the UCT algorithm^50^ balances exploration and exploitation of each parent node to identify desirable mutation nodes. To limit the computational complexity of the search and enable exploration of more nodes, we leverage our proposed efficient multi-mutant strategies described above (e.g., QFF, HPF) after generating possible mutations. The pLM then scores the filtered mutations to quantify the reward for each node. Finally, the mutation node with the highest reward max(*R_m_*) is selected.

### 3.4 Holistic ***in silico*** evaluation framework for protein design

When designing functional proteins, three high-level criteria are critical to performance. First, we want the generated protein sequences to be *relevant* to the design objective, i.e., carry out the function of interest. Since function is primarily encoded via the tertiary structure of the protein, the metrics we devised to assess functional relevance are structure-based. Second, the *quality* of the generated protein sequences can be assessed by their quality to *properly execute* the specified function, i.e., have high fitness^43^. Fitness prediction models have been shown to exhibit relatively high correlation with hundreds of experimental assays^51^ and may therefore be used as an in-silico oracle for true fitness. While these in-silico predictions are imperfect oracles, it should be noted that experiments themselves are imperfect and noisy measurements of the true fitness of proteins^52^. To that end, we leveraged two distinct highly performing zero-shot fitness predictors, such as ESM-1v^9^ (masked language model) and Progen2^19^ (autoregressive model). These independent protein language models were chosen given their complementary inductive biases and strong fitness prediction performance^51^. Lastly, the generated sequences should be *diverse*, i.e., differ from the template wild-type sequence or known natural sequences. We refer to these three design objectives as Relevance, Quality, and Diversity respectively, and crafted two separate metrics for each category (Figure 2). We define these different evaluation metrics as follows:

- **TM-score:** Assesses topological similarity between the predicted structure of the generated sequences and the structure of the template wild type sequence^53^. The final score is the average of the pairwise TM-scores. Structures were predicted with ESMFold^7^. Ideally, the structure of generated sequences is as close to that of the template sequence as possible to ensure functional relevance.
- **pLDDT:** Quantifies the confidence of a structure prediction model (here ESMFold^7^), averaged over the entire sequence. Higher pLDDT has been empirically observed to correlate with higher functional designs^54–56^. Note that this may be limiting in certain cases (e.g. disordered proteins).
- **Overall Fitness:** Reports the proportion of designs with fitness greater than the 75^th^ percentile of fitness scores for natural sequence in the corresponding MSAs to the total number of designs.
- **Maximum Fitness:** Evaluates the ability of the sampling procedure to generate sequences that significantly enhance the fitness of the starting wild-type sequence for the corresponding protein family. For the two aforementioned fitness oracles (ESM-1v and ProGen2) and for each sampled sequence, we compute the ratio of the predicted fitness over that of the wild type sequence. The maximum ratio across designs is then extracted for each predictor and averaged across them.
- **Fréchet ESM Distance (FED):** Assesses the extent to which the generated sequences are distant in embedding space to known natural sequences (e.g., sequences in an MSA for that protein family). It can be seen as a protein-related analog of the Fréchet Inception Distance (FID), initially introduced to evaluate generative adversarial network (GAN)-generated images^57–59^, it has since been used for protein design^40,60^. It leverages the embeddings from the penultimate layer of ESM-1v^9^, recapitulating on broader protein features instead of scores, and computes the Fréchet distance between the embeddings of the generated sequences and the sequences in an MSA for that protein family. It also has a linear correlation to Sequence Dissimilarity (SD), as seen in Table S6-Table S12.
- **Uniqueness:** Reports the proportion of distinct sequences generated. Calculated by taking the ratio of unique sequences to the total number of sequences.
- **Overall Score:** Comprehensively summarises the evaluation by considering metrics from each objective. Computed by taking unique sequences with both a TM-score above 0.5 and fitness above 75^th^ percentile relative to wild-type MSA, averaged across the two fitness oracles involved. We then divide this number by the total number of generated sequences to obtain a score between 0 (bad sampling) and 100 (good sampling).

**Figure 2:**
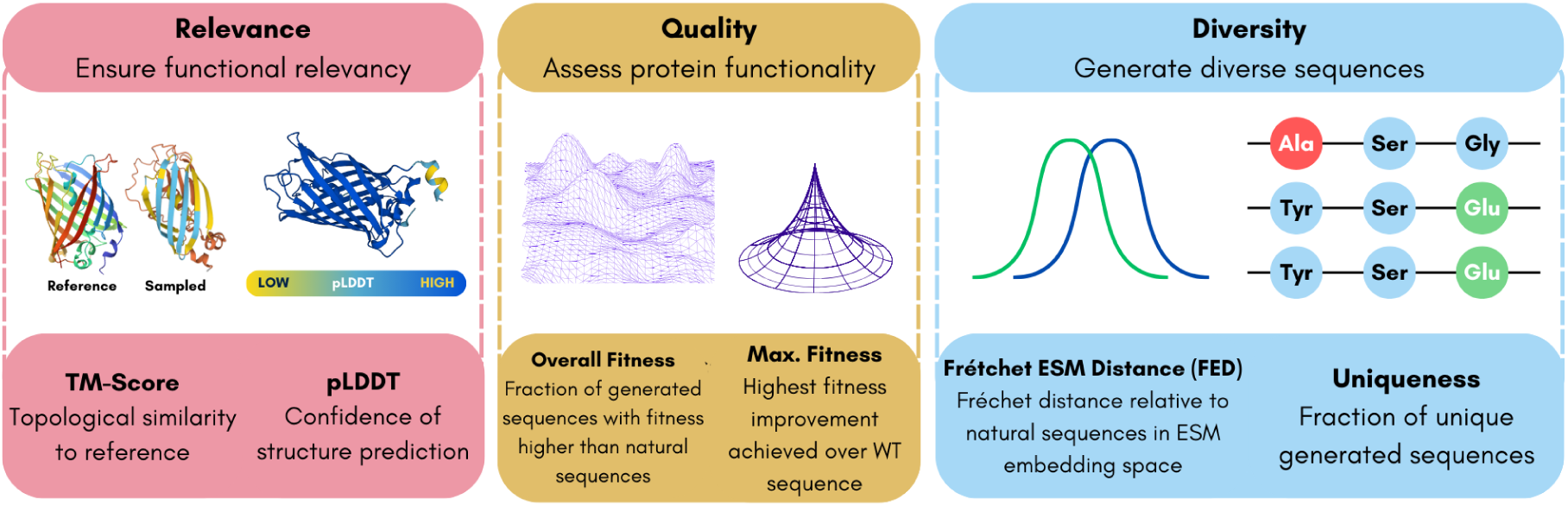
Our holistic *in silico* evaluation framework for functional protein design revolves around three core performance criteria: relevance, quality, and diversity. For each criterion, two complementary metrics enable a thorough comparison of the relative benefits of the various protein language model sampling methods.

### 3.5 Experimental protocol

As we need an autoregressive model that can support both ARS and IRS strategies, we chose Tranception^34^, RITA^18^, and Prot-XLNet^29^ to carry out our experiments as it achieves state-of-the-art fitness prediction performance compared with other autoregressive architectures^51^. For each sampling method, hyperparameter value, and protein family, we ultimately generate 1000 novel designs after several rounds of in silico optimization. Evolution cycles for IRS are set at 4 while the desired length for ARS is equal to the length of the protein wild-type, ensuring all approaches have roughly the same compute budget. Details on datasets and sampling hyperparameters are available in Appendix A.

### 3.6 Selected protein families

To test the generalizability of our results we conducted our experiments for three diverse protein families: *Aequorea victoria* Green Fluorescence Protein (avGFP) (§ B.1), *Escherichia coli* Beta-lactamase TEM-1 (BLAT) (§ B.2), and the Mpro, also known as 3CLpro, region of replicase protein 1ab (R1AB) from the SARS-CoV-2 virus (§ B.3). These proteins represent a diverse selection of proteins and functions, originating from different biological domains — Eukaryote (jellyfish), Prokaryote (E. coli), and Virus (SARS-Cov-2), respectively. They are all well-studied proteins (e.g., all in Swissprot, ProteinGym), well-represented in protein databases (e.g., UniRef) with many natural homologs, and have well-determined tertiary structures. We performed all IRS experiments starting from the corresponding full-length wild-type sequences and used conditioning prompts of at least 5% and up to 85% of the full sequence length for the ARS experiments (Figure S2). Wild-type sequences and MSAs for the 3 protein families were obtained from ProteinGym benchmarks^51^.

## 4 RESULTS AND DISCUSSION

In the experimental evaluation of the methods introduced in § 3, we seek to address the following questions: (1) What are the relative strengths of the various sampling strategies introduced, and their impact on the properties of the generated sequences? (2) What are the benefits of multi-mutant strategies over single-mutant strategies? (3) What are the effects of different sampling hyperparameters on the generated proteins?

### 4.1 Effects of broad sampling approaches

We evaluated the properties of protein generation using both Autoregressive Sampling (ARS) and Iterative Redesign Sampling (IRS) across all categories in our evaluation framework. The best outcomes based on overall score across hyperparameters tested for each sampling method and averaged across protein families, are reported in Figure 3a. We adjust parameters across experiments to ensure a comparable compute budget is used across sampling strategies. We observe that the IRS-doubles approach outperforms *in aggregate* all other sampling strategies. However, while IRS strategies tend to produce generally more relevant and functional samples in aggregate compared with ARS methods, the latter enable the design of a few highly performing designs (i.e., significantly higher maximum fitness value). As proteins generated through IRS method only mutate residues at certain positions, the generated proteins tend to be more similar to the starting wild type sequences (e.g., higher TM-score). Conversely, by providing a fraction of the whole template sequence as prompt, ARS approaches have more flexibility to generate more diverse proteins, at the cost of relevance or quality. The relationship between quality and diversity is shown in Figure 3b for IRS and ARS sampling methods. Additionally, we collected sequences generated through MCMC-based sampling in Biswas et al. ^37^ as additional baseline, focusing our analyses on the 2 protein families covered in that work (presented in Table S6). We observe that MCMC-based sampling in Biswas et al. ^37^ underperforms the various ARS and IRS sampling strategies introduced above.

**Figure 3:**
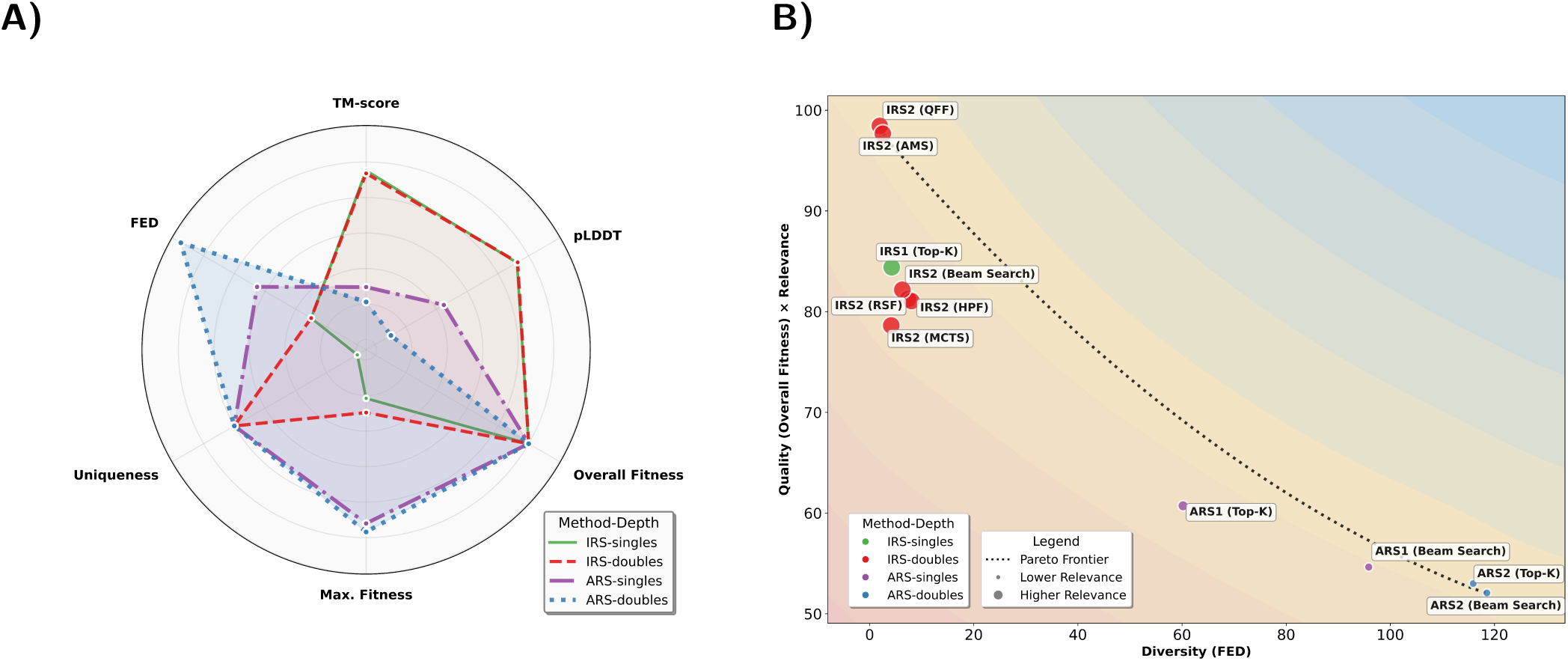
Performance of sampling strategies on our holistic evaluation benchmark. (A) Aggregate performance for IRS and ARS methods. For each category, we select the best combination of sampling method and hyperparameter and report its average performance across the 3 protein families (see detailed results in Table S1). (B) Pareto frontier of sampling performance, highlighting trade-offs between quality, relevance, and diversity metrics for the various sampling strategies.

### 4.2 Benefits of multi-mutant strategies

A comparative summary of the best sampling method and hyperparameter combinations for IRS-singles (single mutation per iteration) and IRS-doubles (double mutations per iteration) based on their overall scores, incorporating the strategies HPF, QFF, AMS, RSF, beam search, and MCTS, is shown in Table S2. In general, IRS-doubles consistently outperform IRS-singles on all metrics. Detailed results are available in Table S6 and results are shown to be consistent with other model architectures presented in Table S4.

We also performed similar experiments for ARS protein generation to compare between single-mutant (ARS-singles) and double-mutant (ARS-doubles) generation. The sampling methods used for comparison are NLP-sampling strategies discussed in § 2.1 and beam search. The beam search implemented in ARS can generate either single or multiple AA extensions, hence its inclusion in both singles and doubles. Results for the best sampling method and hyperparameter combinations for the ARS-singles and ARS-doubles experiments are selected based on the overall score and presented in Table S3. Detailed results are provided in Table S7 with these results consistent with other model architectures presented in Table S5. Unlike IRS strategies, we do not observe any significant benefit of ARS-doubles approaches over ARS-singles.

Within the IRS approach, there are several key observations to be highlighted. The combination of Beam Search with HPF produced proteins with the highest relevance scores, indicating that this method is ideal for generating proteins with the desired function. Detailed performance breakdown across sampling strategies is provided in Table S6. In contrast, QFF with random sampling and AMS produced novel proteins with the highest general score (also shown in Figure 3b), with relevant generated sequences of high fitness and diversity, scoring highly on the two fitness-related metrics. For discovering new and unexplored protein sequences, almost all IRS-doubles methods were highly effective in generating novel proteins. Based on the overall scores, QFF offers a well-rounded balance across all metrics and can thus be considered the most optimal protein generation method. Based on their specific design objectives, practitioners may choose the most appropriate method.

Furthermore, we find that there are significant performance differences across protein families (see detailed results in Tables S8 to S11). The BLAT protein was the most easily evolvable, leading to the highest maximum fitness level across proteins. However, when double-mutations (IRS-doubles) are introduced, other proteins climbed to higher fitness levels and approached the gains achieved in the BLAT experiments (Table S9 and Table S10). This confirmed that multi-mutant approaches can be useful by detecting epistatic effects and making less myopic decisions about which mutations to apply at each step.

The ARS results show that almost all the proteins generated have fitness over 75^th^ percentile of natural sequences. Adjusting sampling temperature to lower values (e.g. 0.01) for beam search and nucleus sampling results in better evaluation scores, consistent with previous findings^61^. Interestingly, when comparing the ARS-singles and ARS-doubles approaches, transitioning to double mutations lead to worse protein generation in terms of relevance (lower TM score and pLDDT), but with positive results in terms of maximum fitness and diversity in aggregate^62^. When searching for a balance between all the metrics, the ARS-singles approach with Top-k achieves the highest overall score, pLDDT, and Uniqueness. Detailed results across different protein families are provided in Table S12. We observe differences in evolvability across the 3 protein families, with BLAT proving to be the most difficult in achieving significant leaps in maximum fitness.

### 4.3 Effects of different sampling hyperparameters

We first sought to investigate the impact of the conditioning prompt length used in ARS on design performance. To that end, we performed sampling experiments across the 3 protein families using prompt lengths based on 5 percentage values (5%, 20%, 40%, 65%, and 85%). Results are reported in Figure S2 and Figure S3. In general, metrics including TM-score, Uniqueness, FED, and Overall Score generally increase as prompt length increases. On the other hand, both pLDDT and maximum fitness exhibit the opposite trend, with 65% length showing the best balance for all the metrics. The impact of prompt length varies across protein families: avGFP and BLAT show moderate changes, while R1AB exhibits drastic increases (FED) or declines (uniqueness and maximum fitness). Longer prompts typically allow ARS methods to generate more family-relevant samples, at the cost of no diversity in the prompting region. Interestingly, diversity also appears to increase with prompt length (up to a limit) as short prompts often lead to degenerate sequences, which fail in relatively similar patterns (e.g., excessive repeat of the same amino acid). We also note that for shorter prompt lengths, the model may generate novel sequences with high predicted fitness but corresponding to a different function than the target, as evidenced by the lower relevance score obtained for these designs. All our subsequent ARS experiments were done with a 65% prompt length for a balanced generation process.

We then aimed to analyze how different hyperparameter values for various sampling methods affect design performance. The hyperparameters we focused on were the values of *k* for Top-k sampling, *p* (or nucleus) for Top-p (nucleus) sampling, *τ* (tau) for typical sampling, and modifying the sampling temperature. For beam search, we tested the impact of the maximum length of subsequent mutations (number of steps) and the number of search rounds for MCTS. To analyze the impact of different hyperparameters on design outcomes, we used Top-k sampling from IRS-singles generation, across the three protein families. The corresponding results are reported in Figure S4. Maximum fitness remained high across all sampling values. However, by prioritizing high-fitness mutations with IRS (i.e. smaller *k* values), proteins generated were more likely to stay within the same family as limited mutations were introduced. Although it provided higher overall fitness, lowering the *k*hyperparameter value led to repetitive mutations in successive generations (i.e. fewer unique proteins generated). Conversely, by increasing the value of *k*, the distance (i.e. FED) between the generated proteins and the starting sequence increased as more diverse mutations were introduced.

## 5 CONCLUSION

In this work, we systematically evaluated the effects of different sampling strategies on the quality, diversity, and relevance of designed protein sequences in silico. We observed a general trade-off between the overall fitness and diversity of the proteins produce across methods: sampling methods that produce greater novelty and diversity in sequences yield proteins that are less functional on average. Selecting the appropriate sampling strategy and fine-tuning hyperparameters is thus crucial for striking the right balance between these competing objectives. Our research also highlights the benefits resulting from developing custom sampling methodologies tailored specifically for protein design, leading to higher aggregate performance across design objectives. Lastly, the observation that the optimal sampling strategy varies across protein family suggests that leveraging several sampling strategies simultaneously could be beneficial. This combined approach would help generate more diverse samples and improve the likelihood of discovering designs with maximum fitness. As the field progressively moves towards developing multi-modal protein language models combining sequence, structure and functional annotations, we anticipate that novel sampling approaches will be needed to fully take advantage of the dependencies across modalities.

## RESOURCE AVAILABILITY

### Lead contact

Requests for further information and resources should be directed to and will be fulfilled by the lead contact upon reasonable request, Jeremie Theddy Darmawan.

### Materials availability

This study did not generate new materials.

### Data and code availability

- We open-source our codebase to provide convenient access to our *in silico* generation and evaluation framework, including all sampling methods, in a unified interface: https://github.com/ i3LBI19-OATML/sampling_plm.
- Any additional information reported in this paper is available from the lead contact upon request.

## ACKNOWLEDGMENTS

- P.N. is supported by GSK and the UK Engineering and Physical Sciences Research Council (EPSRC ICASE award no. 18000077)
- Y.G. holds a Turing AI Fellowship (Phase 1) at the Alan Turing Institute, which is supported by EPSRC grant reference V030302/1.

## AUTHOR CONTRIBUTIONS

Conceptualization, P.N. and J.T.D; methodology, J.T.D and P.N.; investigation, J.T.D and P.N.; writing–original draft, J.T.D and P.N.; writing–review & editing, J.T.D, P.N., and Y.G.; resources, Y.G. and P.N.; supervision, P.N. and Y.G.

## SUPPLEMENTAL INFORMATION

### A Experimental Details

#### A.1 Evaluation Protocol

We applied the relevance, quality, and diversity evaluations contained in our *in silico* evaluation framework discussed in § 3.4 to provide us with the properties of the proteins generated. Both target and reference sequences in FASTA format are required. Target and reference sequences correspond to the generated sequences and naturally occurring sequences, respectively.

#### A.2 Data

Generation using IRS and ARS methods requires an initial protein sequence with a certain length. EVmutation library initiation requires wild-type DMS data from the protein of interest. Proteins that have extensive DMS reports are valid options, including the ones in the ProteinGym benchmarks^51^. Specifically for IRS generation, AA residue PoI is also required for masking and mutation targets.

#### A.3 Sampling and Hyperparameters

Sampling methods that are part of this experiment include random, greedy, Top-k, Top-p, mirostat, and typical sampling as well as beam search and MCTS. Hyperparameters for each are as follows: Mirostat sampling at the best-reported threshold of 3.0^24^, Top-p sampling from 0.1 to 1.0, Typical sampling from 0.1 to 0.95, and Top-k sampling with k values of 2, 3, 5, 10, 15, 20, 30, 40, and 50. Hyperparameter values for beam search and MCTS are 2 and 3.

#### A.4 Prompt Lengths

To address the limitation of shorter prompt lengths, we note that one could alternatively condition on residues starting from the N-terminus and autoregressively generate towards the C-terminus, and then condition on the C-terminus residues and autoregressively generate residues towards the N-terminus. This is possible with models such as Tranception which have been trained to reconstruct sequences autoregressively from both directions.

### B Glossary

#### B.1 avGFP Protein

The jellyfish *Aequorea victoria* has a particular protein that exhibits a green fluorescent when exposed to light^63^. Our experiments were conducted using the *Aequorea victoria* green fluorescent protein (avGFP) data by Sarkisyan et al. ^64^. It is also one of the widely used biophysical property benchmarks and used in other protein generation attempts by Biswas et al. ^37^. Several PoI (66, 65, 148, 203, 205, 145) were identified to be highly determinant for its function^65^.

#### B.2 BLAT Protein

A fourth of the proteins in *Escherichia coli* are enzymes and beta-lactamase TEM-1 is an enzyme that confers resistance to the bacteria^66^. Resistance is conferred by hydrolyzing *β*-lactam rings of penicillins and other common antibiotics found in healthcare facilities^67^. It also has a well-studied 3D structure and thermodynamic range that allow it to be suitable for medical investigations into drug resistance. The PoI at 36, 164, 179, 182, and 250 are critical for executing its function^66,68^.

#### B.3 R1AB Protein

Belonging to the SARS-CoV-2 virus, the Mpro region of the R1AB protein (also known as the main protease Mpro) and its substrates play an essential functional role throughout every step of the viral life cycle. The protein is a peptidase that initiates autoproteolysis of the pp1a and pp1ab polyprotein translated from the ORF1 gene which is essential for viral replication^69^. Residues at locations 145, 41, 299, 142, and 189 are PoI that were identified by Flynn et al. ^69^ that significantly contribute to its proteolytic activity.

#### B.4 Top-k Sampling

The sorting of probabilities (or scores) in descending order and zero-masking the values that are beyond the *k* rank. It truncates unreliable, lower probability tokens and considers only the highest *k* probable tokens.

#### B.5 Greedy Sampling

A variant of Top-k sampling that only considers the highest (top-1) probable token.

#### B.6 Beam Search

Beam search is a greedy, heuristic search algorithm that looks ahead into the next *N* tokens. It considers the best combination of all proceeding tokens ahead of the current token. Although computationally costly, this approach would generally generate better results over Top-k sampling as it takes into account the next *N* combinations that might be missed when selecting individual tokens for each position.

#### B.7 Top-p Sampling

As a solution to the Top-k sampling and beam search that is prone to the boredom trap, Top-p sampling (or nucleus sampling) was developed by Holtzman et al. ^22^. It samples from a pool of the smallest possible set of tokens in which the cumulative probability exceeds the probability p, also called the nucleus. Compared to Top-k sampling, this approach has a dynamically adjusting sampling pool according to the predicted probability distribution of the language model.

#### B.8 Mirostat Sampling

Based on reports that text quality is most desirable at certain likelihood ranges, mirostat sampling was developed to keep the generated text within a predetermined perplexity value^24^. Therefore, high-quality text could be obtained without any fine-tuning. This approach is also able to prevent the generated text from the boredom trap (repetitions) as well as the confusion trap (incoherence).

#### B.9 Typical Sampling

Inspired by the psycholinguistic theory of human communication, Meister et al. ^23^ formally defines a criterion for text generation that minimizes the next generated token information content as close as possible to the model’s conditional entropy (or expected information content). Using this information theory foundation, typical sampling efficiently enforces this criterion upon text generation.

#### B.10 Fréchet distance

The objective of generative models is to imitate the original data as best as possible. Hence, the distance between the generated data *p_w_*(.) and *p*(.), or also called the Fréchet distance, can be a measure of generation quality^58,70^. This measure is also known as the Wasserstein-2 distance^71^. The Fréchet distance *d*(.,.) is calculated between the *p*(.) Gaussian with mean (*m, C*) and the *p_w_*(.) Gaussian with mean (*m_w_, C_w_*). Since we only modified the Inception model used in Heusel et al. ^58^ into ESM-1v that is suitable for protein sequences, the “Fréchet Inception Distance” (FID) equation^72^ still applies:

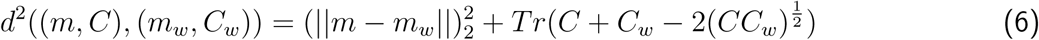

### C EVmutation Initialization

Initializing the EVmutation^49^ model requires two steps: (1) collecting deep multiple sequence alignments (MSA) and (2) generating the model parameters. MSAs were collected from the ProteinGym database for each protein family^51^. We used the recommended plmc^49^ commands to accomplish this with L2-regularization on the couplings set, applying their formula.

### D Figures and Tables

**Figure S1:**
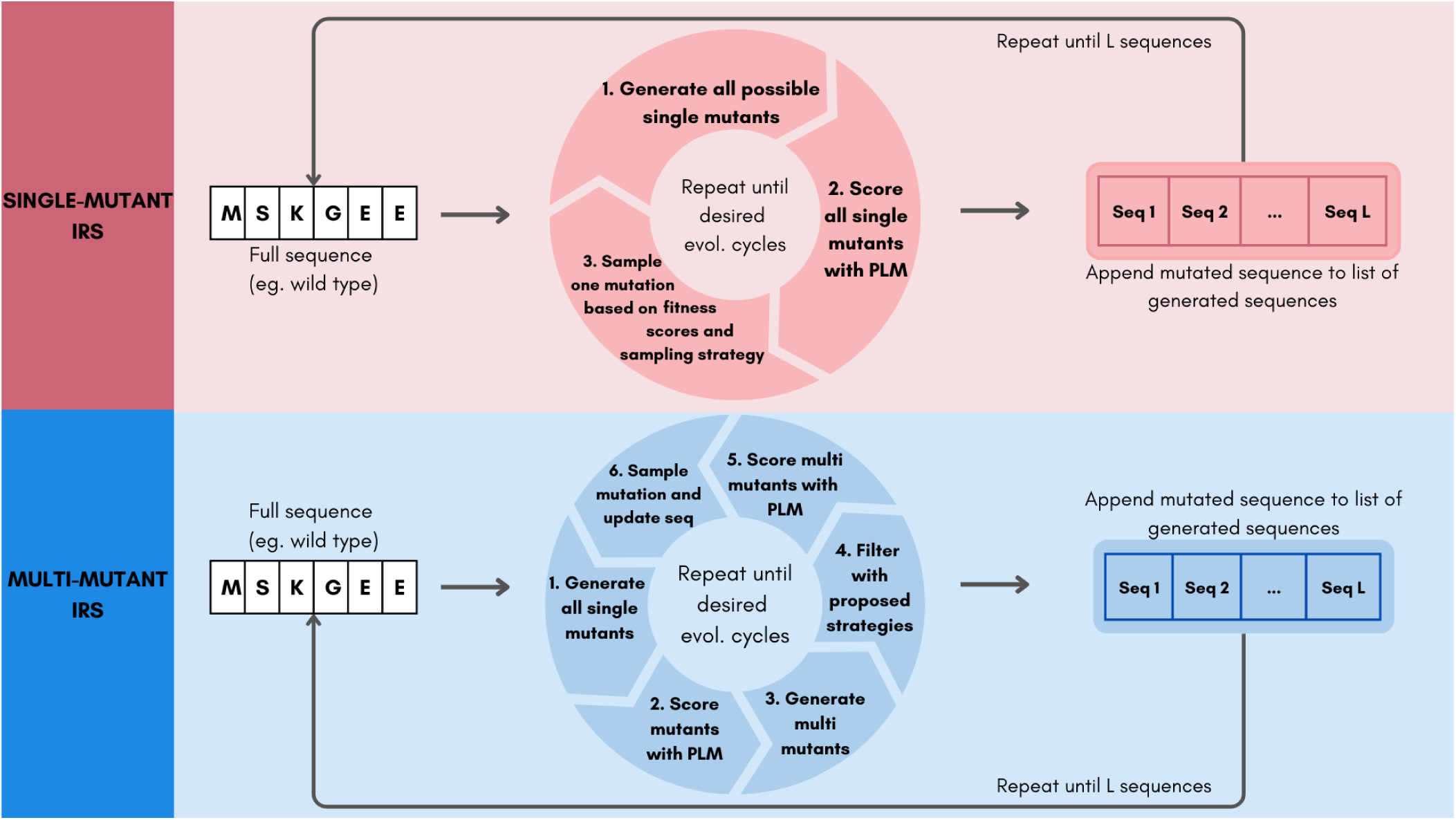
High-level single and multi-mutant IRS algorithms for generating novel protein sequences. Detailed algorithms are provided in supplementary (Algorithm S7 and Algorithm S9).

**Figure S2:**
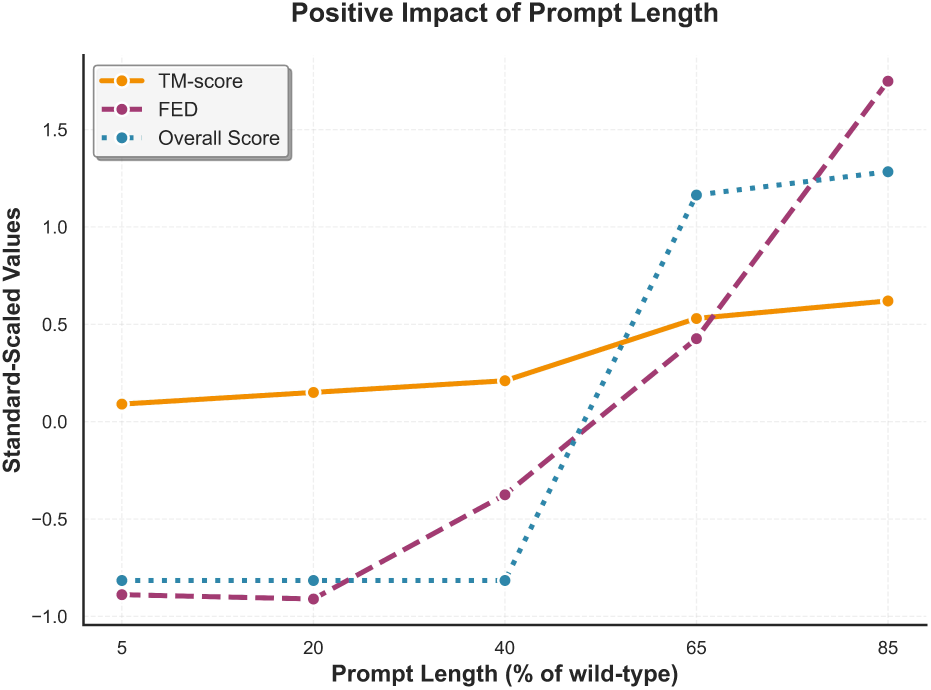
Impact of prompt lengths on sampling quality. Results for a particular prompt length are obtained using the best method and hyperparameter, averaged across families.

**Figure S3:**
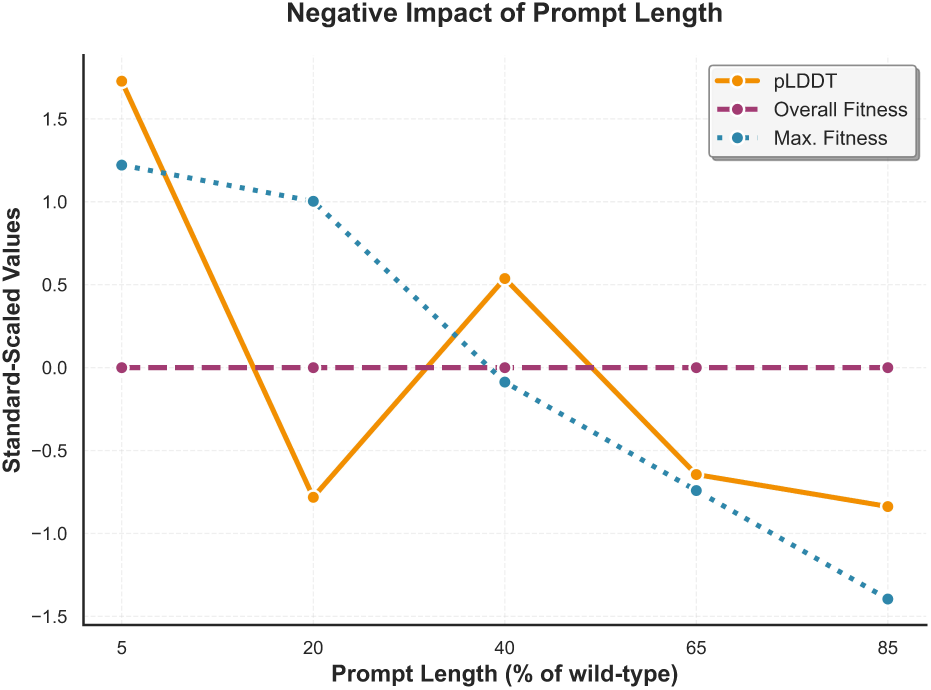
Negative impact of prompt lengths based on percentage of wild-type prompted on our evaluation framework. The values for a particular prompt length were obtained using the best combination across the experimented proteins and models. Standardization was performed to plot each of the effects.

**Figure S4:**
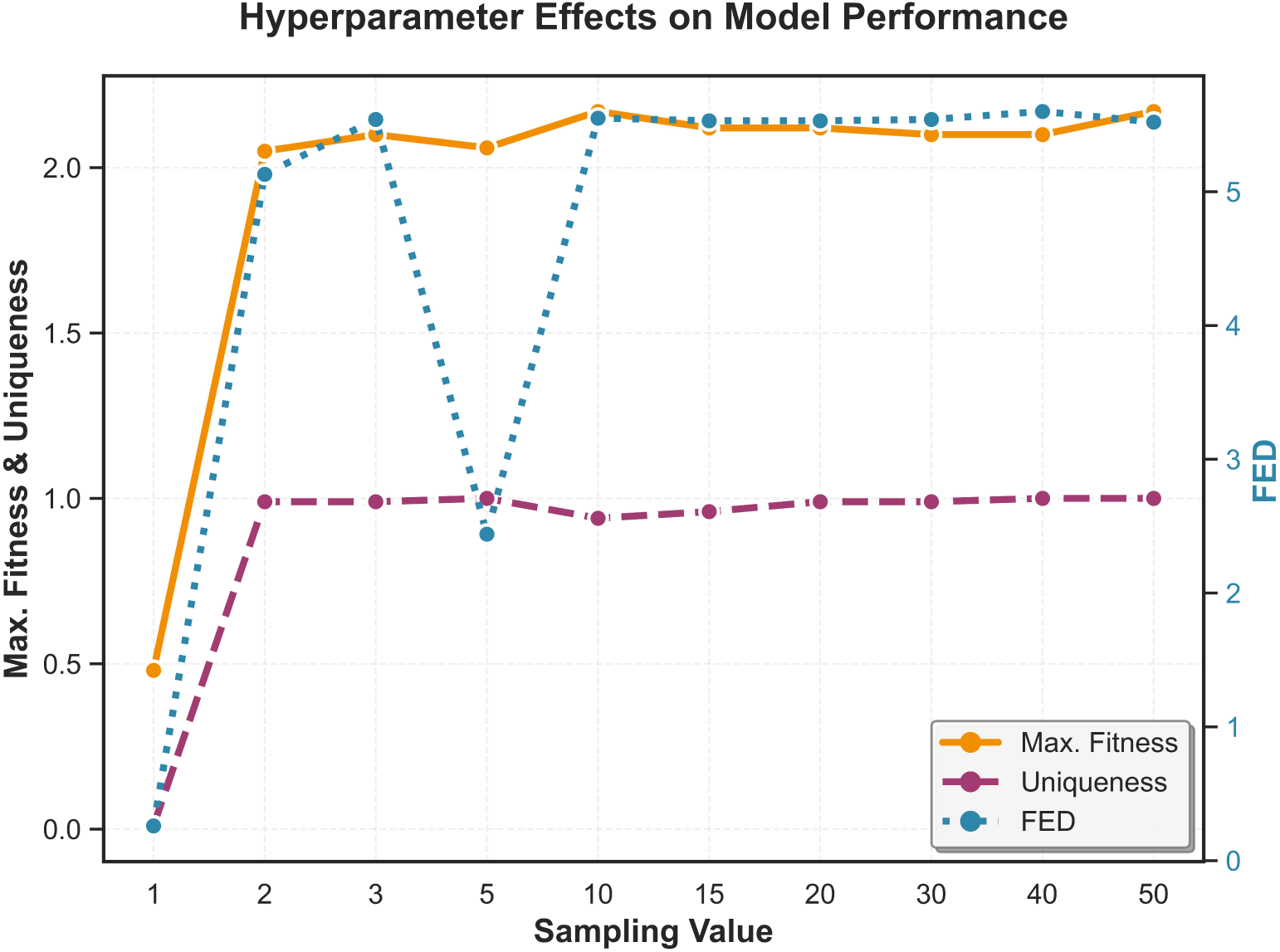
**Comparing the effects of Top-k hyperparameters on Max. Fitness, Uniqueness, and FED metrics.**

**Table S1:** Comparing across ARS and IRS (singles and doubles) methods on the different aspects of the evaluation framework. Bold denotes best scores.

| Method | Relevance | | Quality | | Diversity | | Overall Score ↑ |
| --- | --- | --- | --- | --- | --- | --- | --- |
| | TM-score ↑ | pLDDT ↑ | Overall Fitness ↑ | Max. Fitness ↑ | Uniqueness ↑ | FED ↑ | |
| ARS-singles | 0.60 | 71.94 | **97.50** | 3.76 | **1.00** | 50.22 | 94.00 |
| ARS-doubles | 0.55 | 54.91 | **97.50** | **3.87** | **1.00** | **114.00** | 86.50 |
| IRS-singles | **0.99** | 95.57 | 92.10 | 2.07 | 0.84 | 5.45 | 77.00 |
| IRS-doubles | 0.98 | **95.79** | 97.17 | 2.26 | **1.00** | 5.54 | **97.00** |

**Table S2:** Comparison for IRS strategies. Best combination of sampling method and hyperparameter are reflected, averaged across protein families. Detailed results are in Table S8, Table S9, and Table S10. Other model results are in Table S4. Bold indicates the best scores for each model.

| Depth | Sampling | Relevance | | Quality | | Diversity | | Overall Score ↑ |
| --- | --- | --- | --- | --- | --- | --- | --- | --- |
| | | TM-score ↑ | pLDDT ↑ | Overall Fitness ↑ | Max. Fitness ↑ | Uniqueness ↑ | FED ↑ | |
| Singles | Top-K | **0.99 ± 0.01** | 95.51 ± 0.24 | 84.45 ± 0.94 | 2.13 ± 0.04 | **1.00 ± 0** | 4.26 ± 0 | 84.45 ± 1.91 |
| Doubles | QFF | 0.96 ± 0.01 | **95.91 ± 0.18** | **99.83 ± 2.31** | **2.35 ± 0.13** | **1.00 ± 0** | 2.55 ± 0 | **99.83 ± 1.72** |
| | HPF | 0.98 ± 0.01 | 95.38 ± 0.20 | 81.74 ± 0.96 | 2.06 ± 0.18 | **1.00 ± 0** | 7.39 ± 0 | 81.75 ± 1.87 |
| | AMS | 0.97 ± 0.01 | 95.65 ± 0.21 | 98.67 ± 1.96 | 2.20 ± 0.03 | **1.00 ± 0** | 1.97 ± 0 | 98.67 ± 1.41 |
| | RSF | 0.97 ± 0 | 95.25 ± 0.11 | 82.03 ± 1.80 | 2.04 ± 0.10 | **1.00 ± 0** | **8.06 ± 0** | 82.03 ± 1.34 |
| | MCTS | **0.99 ± 0.00** | 95.62 ± 0.18 | 78.58 ± 2.15 | 2.15 ± 0.06 | **1.00 ± 0** | 6.29 ± 0 | 78.59 ± 1.87 |
| | Beam Search | 0.98 ± 0.01 | 95.53 ± 0.19 | 82.61 ± 0.97 | 2.06 ± 0.11 | **1.00 ± 0** | 4.17 ± 0 | 82.62 ± 1.98 |

**Table S3:** Comparison for ARS strategies. Values reflect the best sampling method and hyperparameter combination. Detailed results are in Table S12. Other model results are in Table S5. Bold indicates the highest scores for each model.

| Depth | Sampling | Relevance | | Quality | | Diversity | | Overall Score ↑ |
| --- | --- | --- | --- | --- | --- | --- | --- | --- |
| | | TM-score ↑ | pLDDT ↑ | Overall Fitness ↑ | Max. Fitness ↑ | Uniqueness ↑ | FED ↑ | |
| Singles | Top-K | **0.55 ± 0.01** | **65.00 ± 0.39** | 98.49 ± 0.18 | 3.87 ± 0.14 | **1.00 ± 0** | 60.21 ± 0 | **81.13 ± 2.31** |
| | Beam Search | 0.52 ± 0.01 | 55.07 ± 0.54 | **99.30 ± 0.12** | **4.90 ± 0.08** | **1.00 ± 0** | 95.86 ± 0 | 80.06 ± 2.41 |
| Doubles | Top-K | 0.53 ± 0.01 | 51.99 ± 0.50 | 98.27 ± 0.70 | 3.31 ± 0.16 | **1.00 ± 0** | 25.30 ± 0 | 79.16 ± 2.29 |
| | Beam Search | 0.53 ± 0.01 | 52.31 ± 0.50 | 96.25 ± 1.03 | 3.21 ± 0.13 | **1.00 ± 0** | **115.93 ± 0** | 77.93 ± 2.34 |

**Table S4:** Complete comparison for IRS strategies across Tranception, RITA, and Prot-XLNet. Best combination of sampling method and hyperparameter are reflected, averaged across protein families. Detailed results for Tranception are in Table S8, Table S9, and Table S10. Bold indicates the best scores for each model.

| Depth | Sampling | Relevance |  | Quality |  | Diversity |  | Overall Score ↑ |
| --- | --- | --- | --- | --- | --- | --- | --- | --- |
|  |  | TM-score ↑ | pLDDT ↑ | Overall Fitness ↑ | Max. Fitness ↑ | Uniqueness ↑ | FED ↑ |  |
| Tranception |  |  |  |  |  |  |  |  |
| Singles | Top-K | 1.00 | 95.88 | 99.82 | 2.30 | 0.98 | 2.75 | 86.07 |
| Doubles | QFF | 0.96 | 95.88 | 100.00 | 2.41 | 0.98 | 1.89 | 100.00 |
|  | HPF | 0.99 | 95.79 | 95.75 | 2.30 | 0.98 | 2.68 | 95.75 |
|  | AMS | 0.99 | 95.74 | 100.00 | 2.36 | 0.98 | 2.10 | 99.10 |
|  | RSF | 0.98 | 95.74 | 95.58 | 2.27 | 0.98 | 2.78 | 95.58 |
|  | MCTS | 0.99 | 95.74 | 91.32 | 2.29 | 0.98 | 2.77 | 91.18 |
|  | Beam Search | 0.99 | 95.82 | 98.36 | 2.30 | 0.98 | 2.66 | 98.27 |
| RITA |  |  |  |  |  |  |  |  |
| Singles | Top-K | 0.99 | 95.88 | 100.00 | 2.20 | 0.98 | 2.84 | 93.00 |
| Doubles | QFF | 0.97 | 96.04 | 100.00 | 2.36 | 0.98 | 1.97 | 100.00 |
|  | HPF | 0.99 | 95.81 | 99.00 | 2.19 | 0.98 | 2.69 | 99.00 |
|  | AMS | 0.99 | 95.97 | 100.00 | 2.14 | 0.98 | 2.81 | 100.00 |
|  | RSF | 0.98 | 95.62 | 100.00 | 2.21 | 0.98 | 2.69 | 100.00 |
|  | MCTS | 0.99 | 95.90 | 90.00 | 2.18 | 0.98 | 2.65 | 90.00 |
|  | Beam Search | 0.99 | 95.84 | 99.50 | 2.26 | 0.98 | 2.63 | 99.50 |
| Prot-XLNet |  |  |  |  |  |  |  |  |
| Singles | Top-K | 0.98 | 94.77 | 53.50 | 1.92 | 0.98 | 7.18 | 53.50 |
| Doubles | QFF | 0.96 | 95.80 | 99.50 | 2.32 | 0.98 | 2.05 | 99.50 |
|  | HPF | 0.97 | 94.54 | 50.50 | 1.81 | 0.98 | 16.81 | 50.50 |
|  | AMS | 0.93 | 95.25 | 96.00 | 2.14 | 0.98 | 2.73 | 96.00 |
|  | RSF | 0.96 | 94.40 | 50.50 | 1.75 | 0.98 | 18.70 | 50.50 |
|  | MCTS | 0.99 | 95.22 | 54.50 | 2.02 | 0.98 | 7.10 | 54.50 |
|  | Beam Search | 0.97 | 94.93 | 50.00 | 1.69 | 0.98 | 13.59 | 50.00 |

**Table S5:** Complete comparison for ARS strategies across Tranception, RITA, and Prot-XLNet. Values reflect the best sampling method and hyperparameter combination. Detailed results for Tranception are in Table S12. Bold indicates the highest scores for each model.

| Depth | Sampling | Relevance |  | Quality |  | Diversity |  | Overall Score ↑ |
| --- | --- | --- | --- | --- | --- | --- | --- | --- |
|  |  | TM-score ↑ | pLDDT ↑ | Overall Fitness ↑ | Max. Fitness ↑ | Uniqueness ↑ | FED ↑ |  |
| Tranception |  |  |  |  |  |  |  |  |
| Singles | Top-K | 0.65 | 83.63 | 95.96 | 2.19 | 0.98 | 1.32 | 95.96 |
|  | Beam Search | 0.53 | 59.30 | 97.91 | 5.56 | 0.98 | 100.01 | 85.32 |
| Doubles | Top-K | 0.54 | 54.70 | 97.72 | 4.43 | 0.98 | 110.24 | 85.22 |
|  | Beam Search | 0.55 | 54.21 | 93.64 | 3.86 | 0.98 | 117.30 | 83.68 |
| RITA |  |  |  |  |  |  |  |  |
| Singles | Top-K | 0.44 | 63.07 | 100.00 | 5.70 | 0.98 | 69.11 | 60.00 |
|  | Beam Search | 0.49 | 56.89 | 100.00 | 5.32 | 0.98 | 81.87 | 71.00 |
| Doubles | Top-K | 0.49 | 53.63 | 99.50 | 3.30 | 0.98 | 107.76 | 68.50 |
|  | Beam Search | 0.48 | 54.55 | 99.00 | 4.09 | 0.98 | 105.21 | 67.00 |
| Prot-XLNet |  |  |  |  |  |  |  |  |
| Singles | Top-K | 0.56 | 48.30 | 99.50 | 4.05 | 0.98 | 110.21 | 87.50 |
|  | Beam Search | 0.53 | 49.04 | 100.00 | 4.02 | 0.98 | 105.69 | 84.00 |
| Doubles | Top-K | 0.55 | 47.64 | 97.50 | 2.56 | 0.98 | 129.78 | 84.00 |
|  | Beam Search | 0.56 | 48.20 | 96.00 | 1.96 | 0.98 | 133.14 | 83.00 |

**Table S6:**
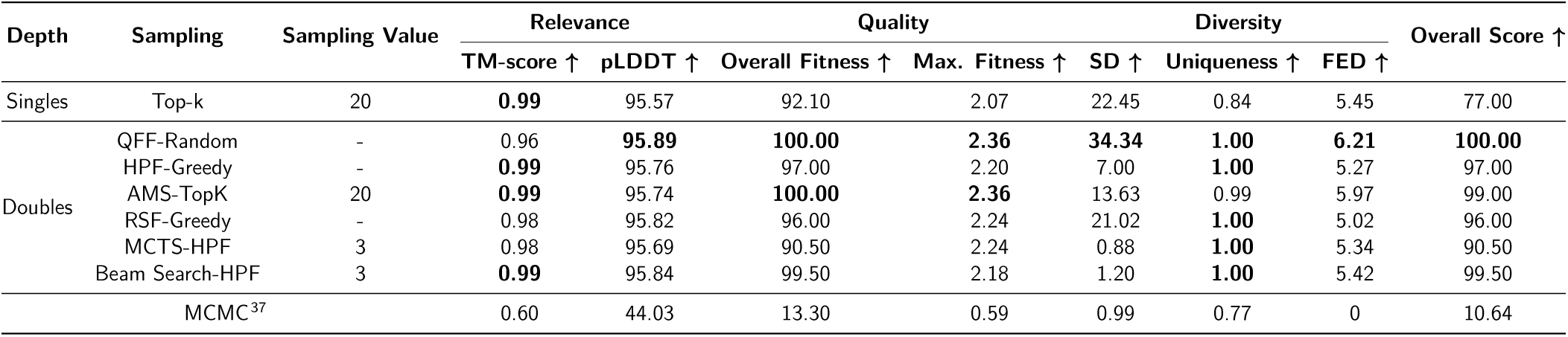
Detailed comparison between the IRS protein generation classes for protein family: avGFP and BLAT. Detailed best results for each method are available in Table S8 (IRS) and Table S9 and Table S10 (IRS-doubles). Bold denotes best scores. Results here are identical as with the inclusion of 3 protein families in Table S2. Sampling value refers to the value of the main sampling hyperparameter for a given method (i.e., k for top-k, p for top-p, etc.).

**Table S7:**
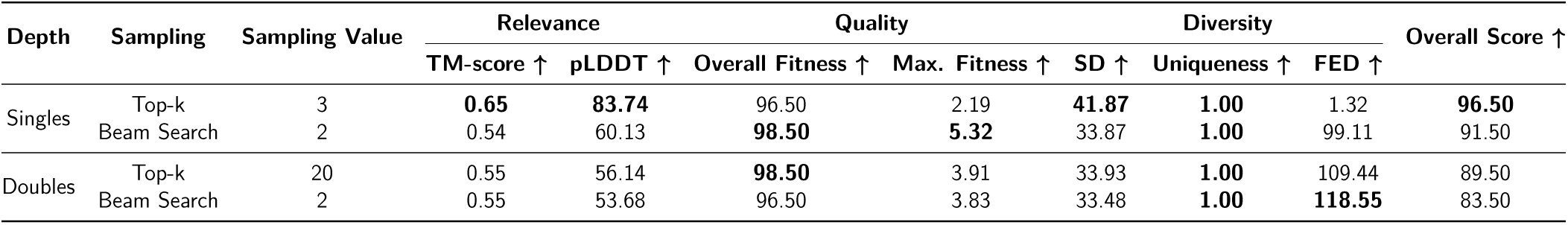
Detailed comparison between the ARS protein generation class. Detailed best results for each method are available in Table S12. Bold denotes best scores. Sampling value refers to the value of the main sampling hyperparameter for a given method (i.e., k for top-k, p for top-p, etc.).

**Table S8:**
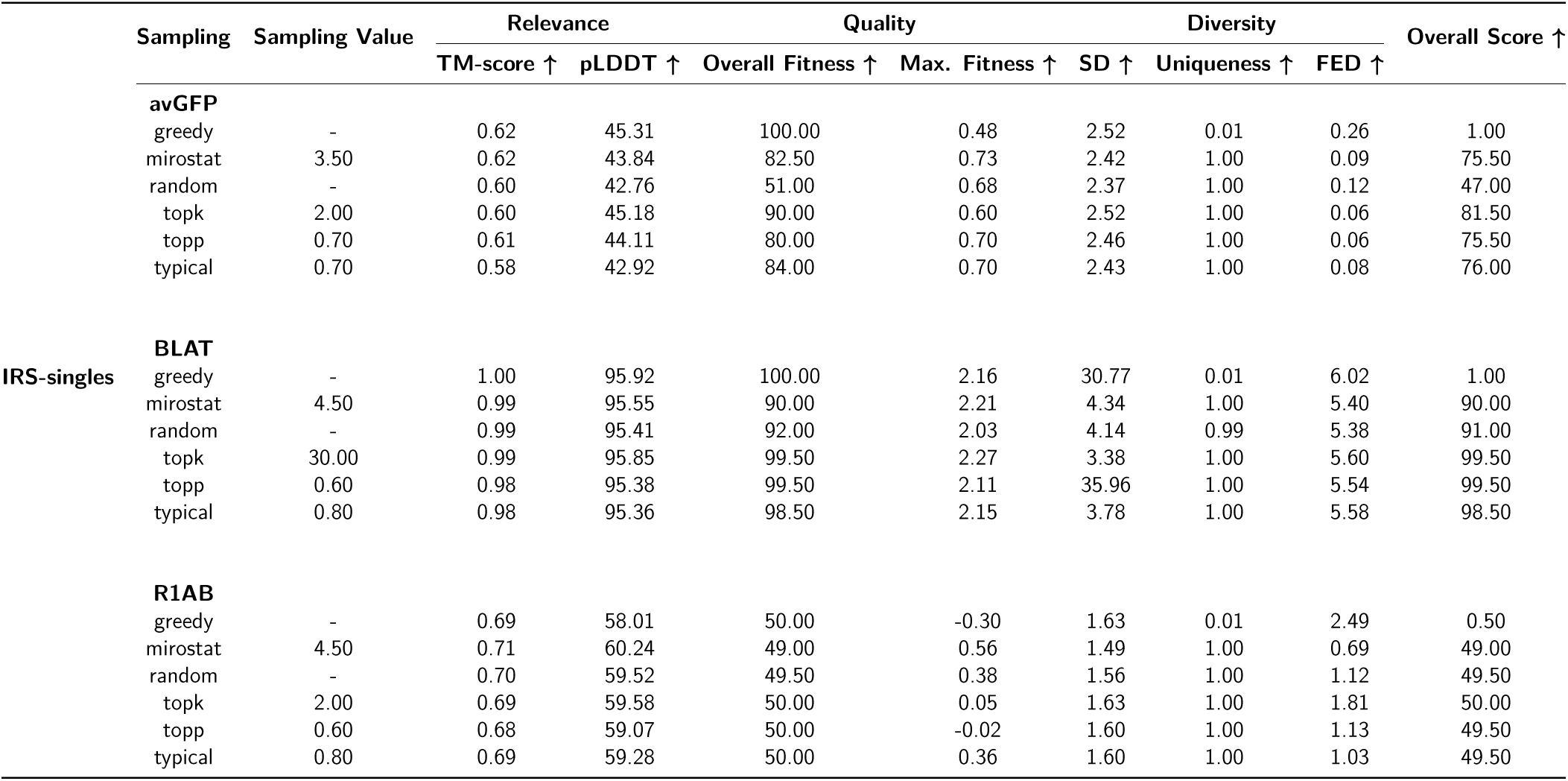
Performance of the best sampling methods and hyperparameter values on IRS-singles protein generation across 3 proteins experimented. Sampling value refers to the value of the main sampling hyperparameter for a given method (i.e., k for top-k, p for top-p, etc.).

**Table S9:**
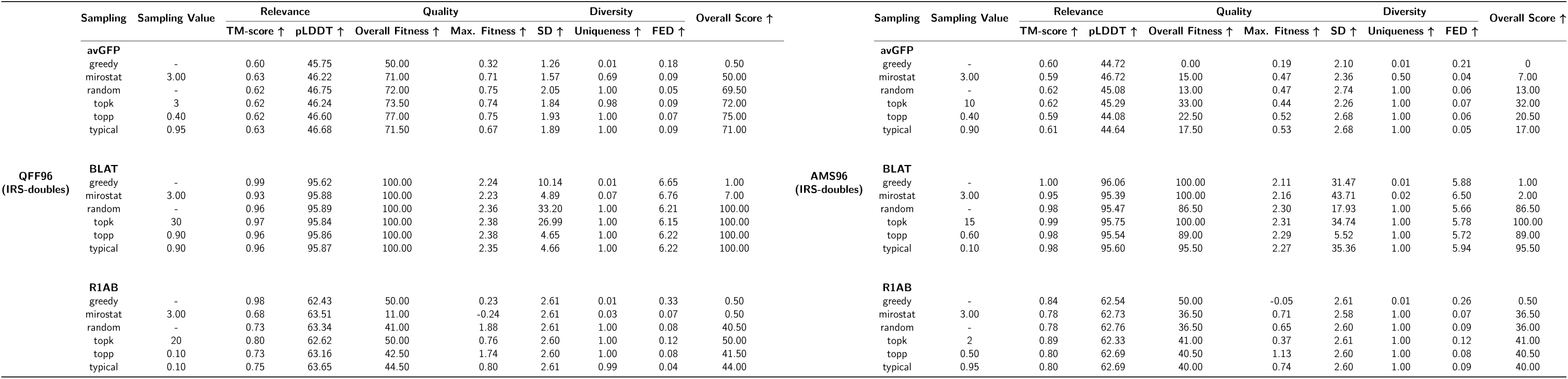
Performance of the best sampling methods and hyperparameter values on IRS-doubles QFF (left) and AMS (right) protein generation across 3 proteins experimented. Sampling value refers to the value of the main sampling hyperparameter for a given method (i.e., k for top-k, p for top-p, etc.).

**Table S10:**
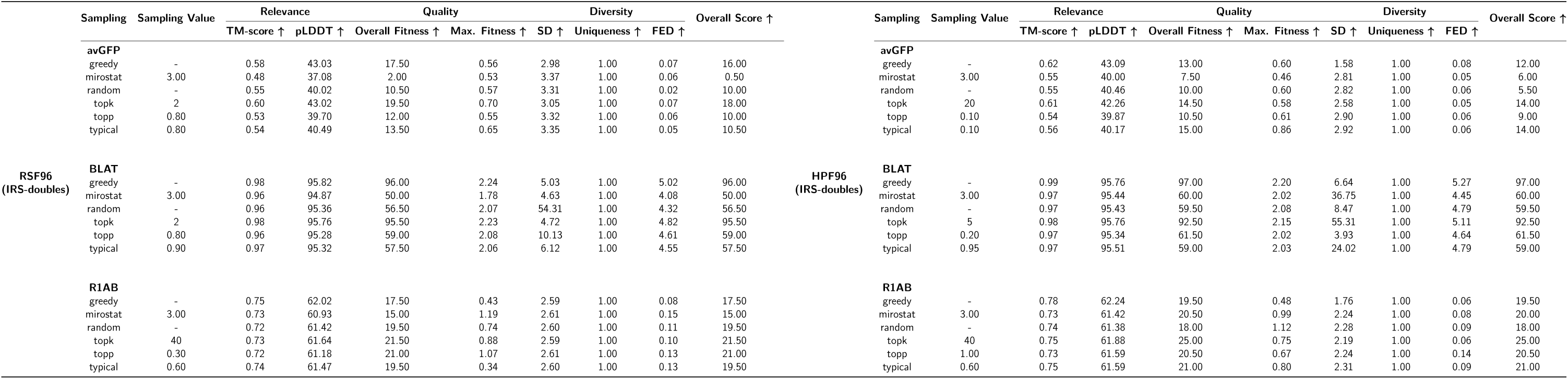
Performance of the best sampling methods and hyperparameter values on IRS-doubles RSF (left) and HPF (right) protein generation across 3 proteins experimented. Sampling value refers to the value of the main sampling hyperparameter for a given method (i.e., k for top-k, p for top-p, etc.).

**Table S11:**
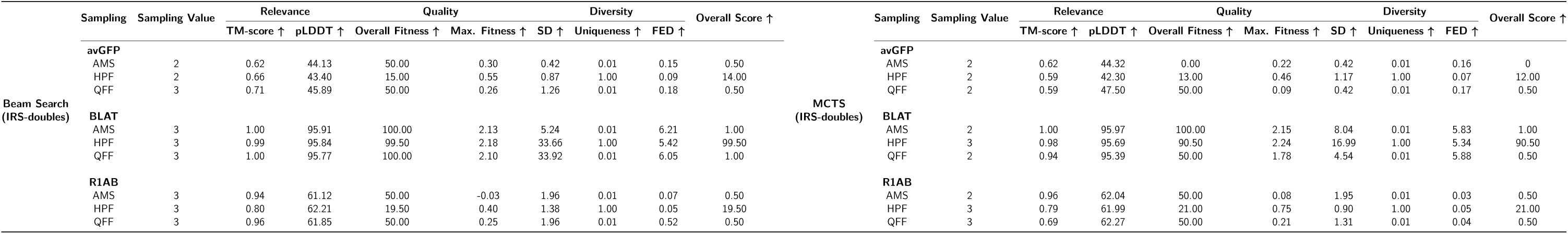
Performance of the best sampling methods and hyperparameter values on IRS-doubles Beam Search and MCTS protein generation across 3 proteins experimented. Sampling value refers to the value of the main sampling hyperparameter for a given method (i.e., k for top-k, p for top-p, etc.).

**Table S12:**
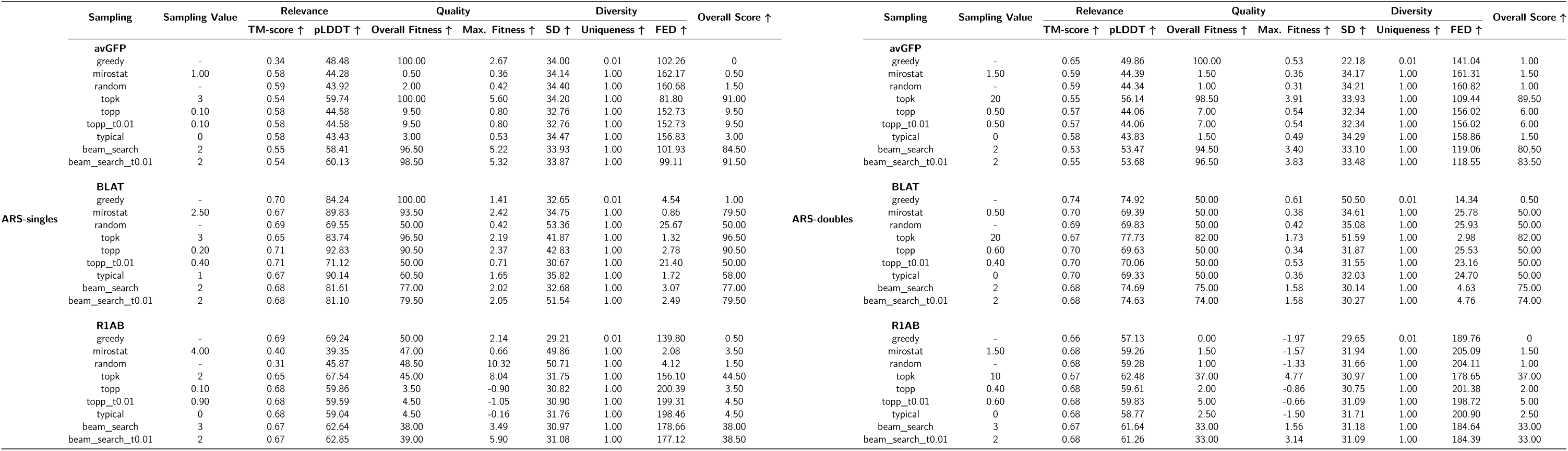
Performance of the best sampling methods and hyperparameter values on ARS-singles and ARS-doubles protein generation across 3 proteins experimented. Sampling value refers to the value of the main sampling hyperparameter for a given method (i.e., k for top-k, p for top-p, etc.).

### E Algorithms

#### Algorithm S1

Mirostat sampling^24^

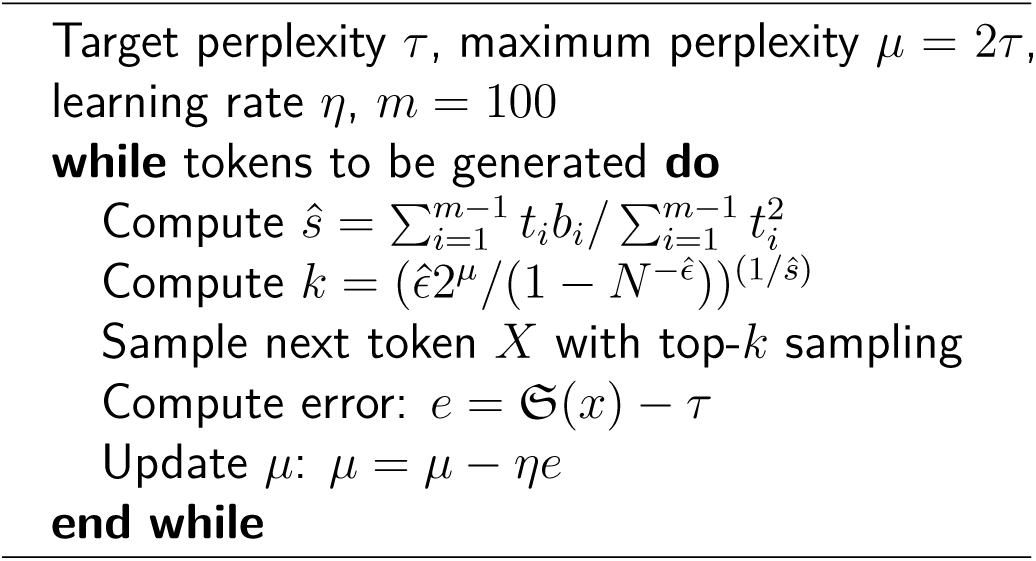

#### Algorithm S2

Quantitative-Function Filter (QFF) pseudocode

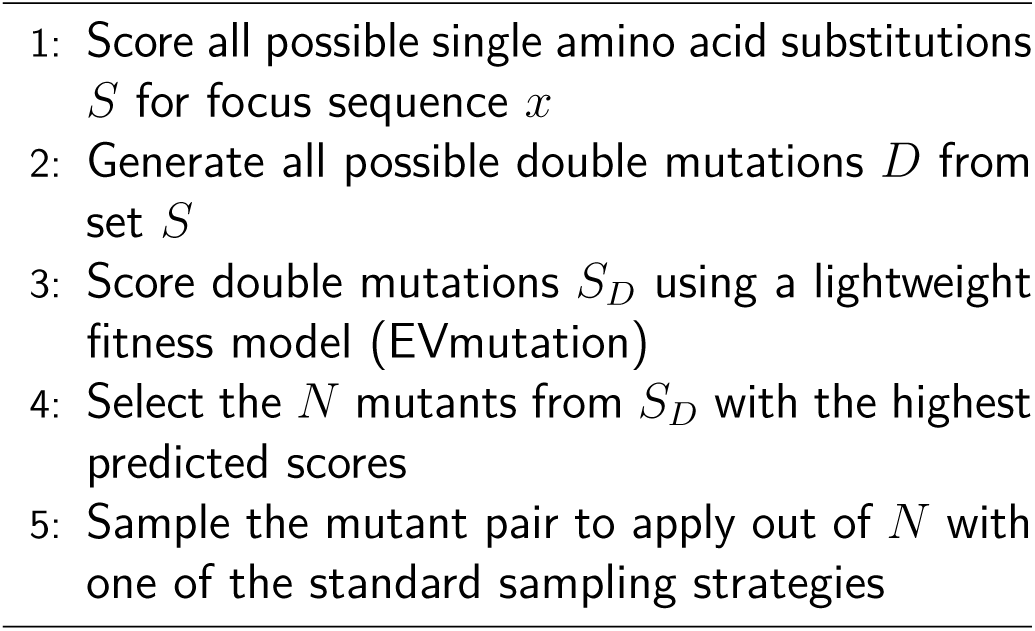

#### Algorithm S3

High-Probability Filter (HPF) pseudocode

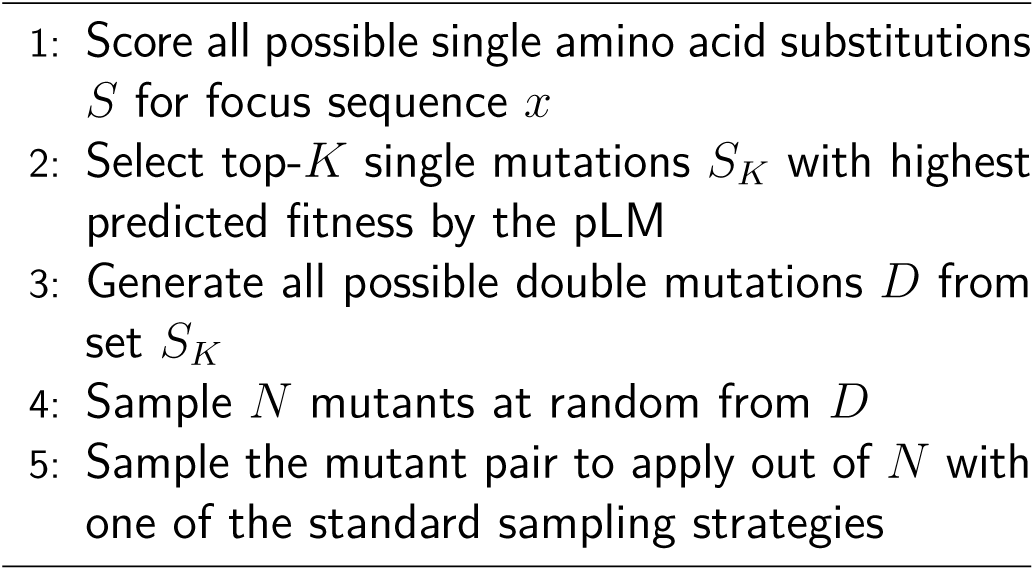

#### Algorithm S4

Attention-Matrix Sampling (AMS) pseudocode

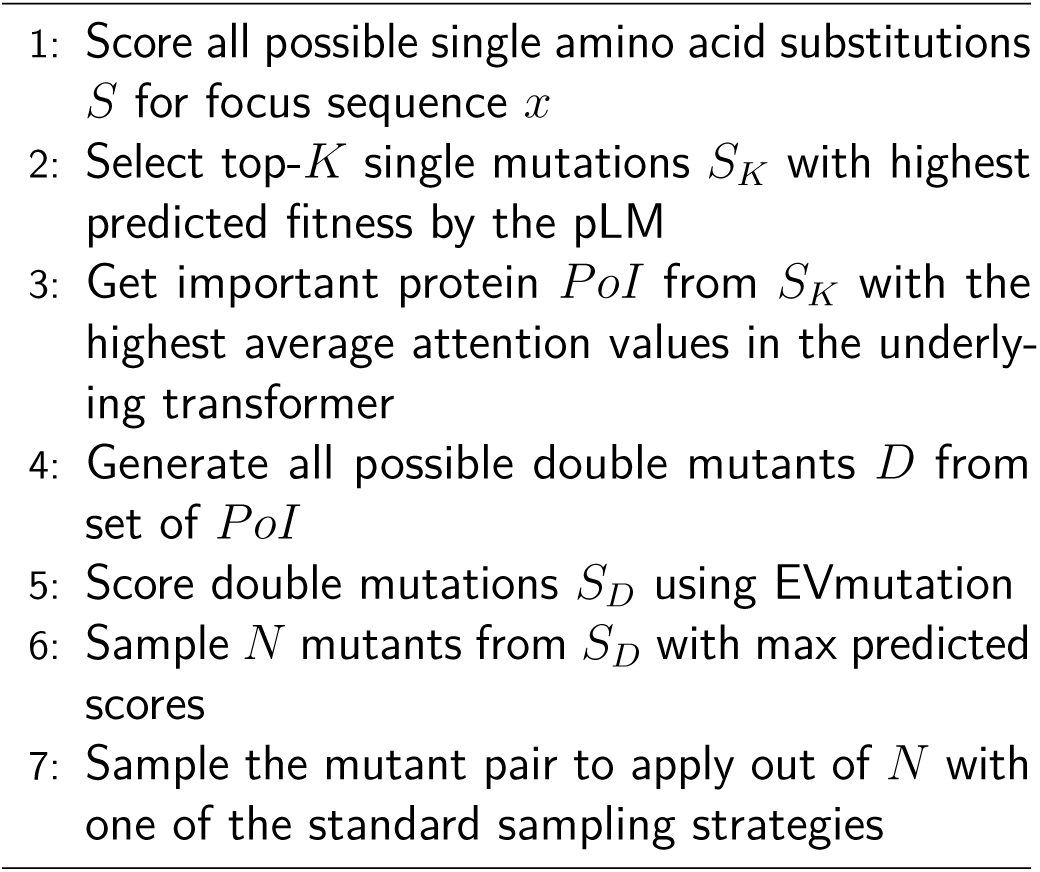

#### Algorithm S5

Random-Stratified Filter (RSF) pseudocode

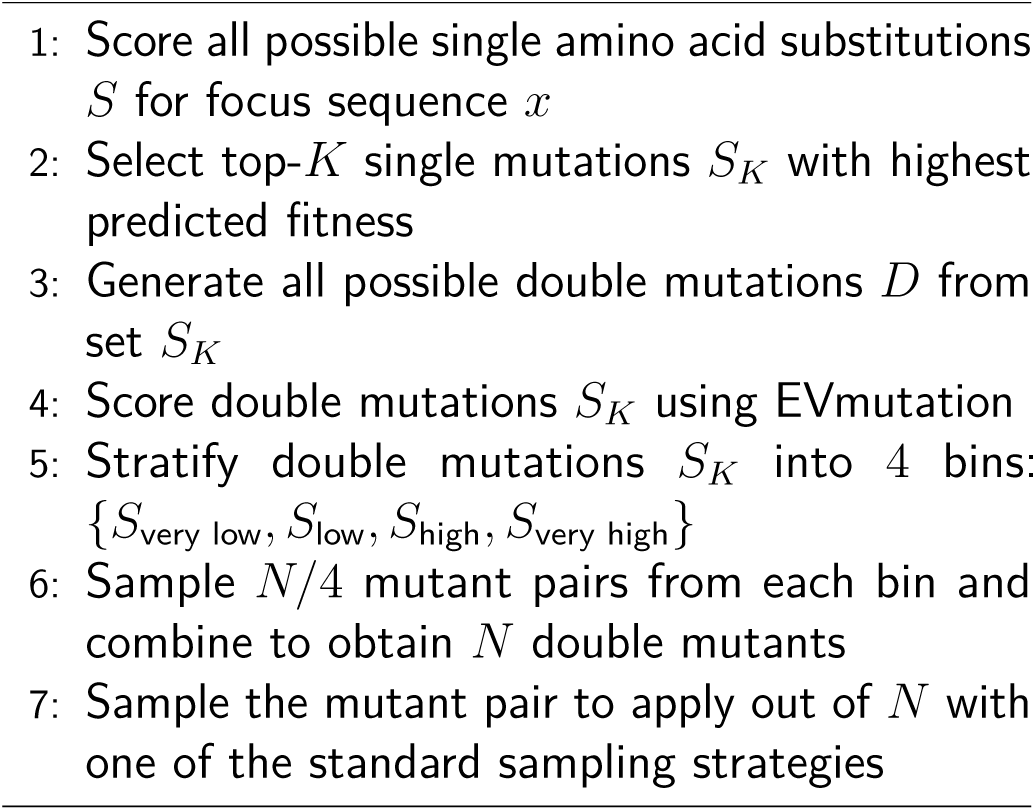

#### Algorithm S6

MCTS Pseudocode

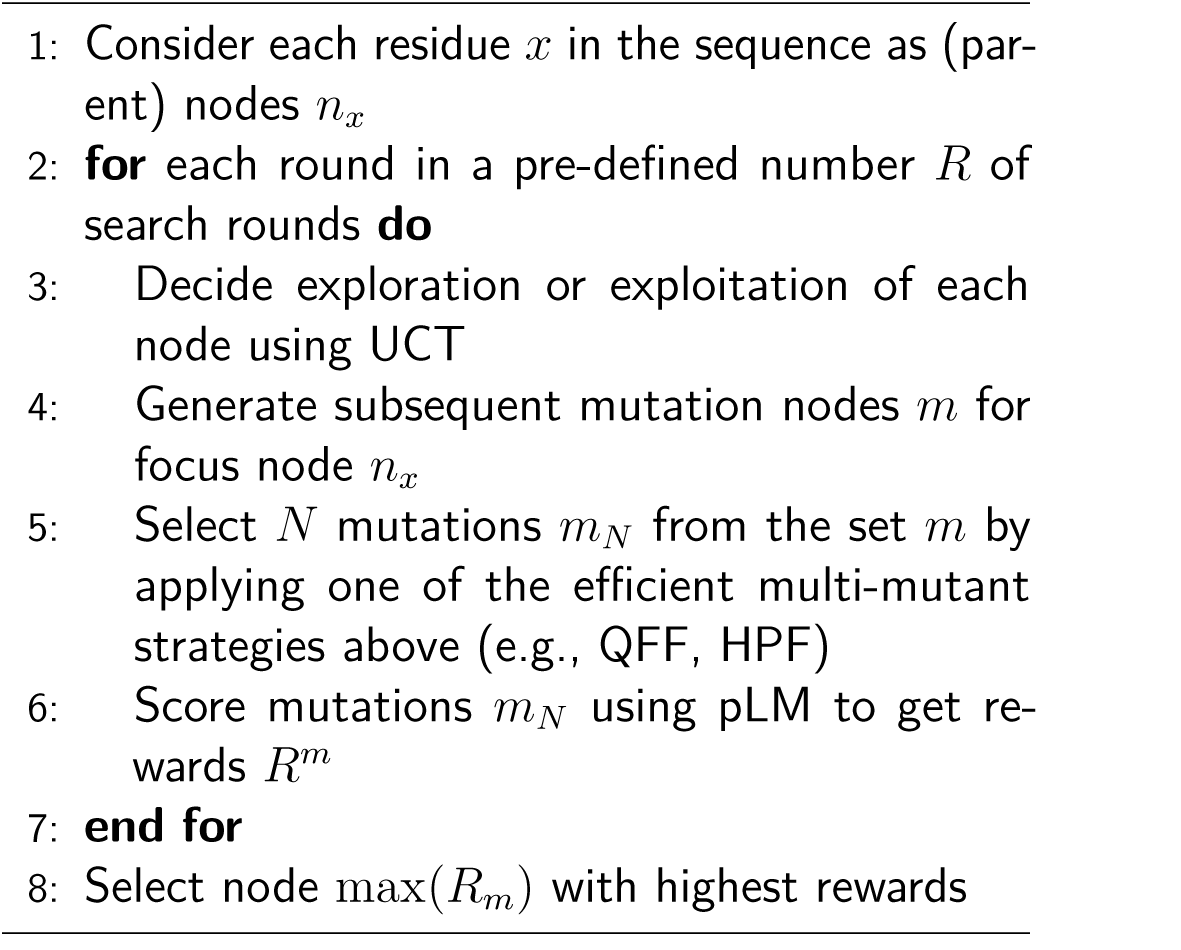

#### Algorithm S7

IRS-singles Protein Generation Pseudocode

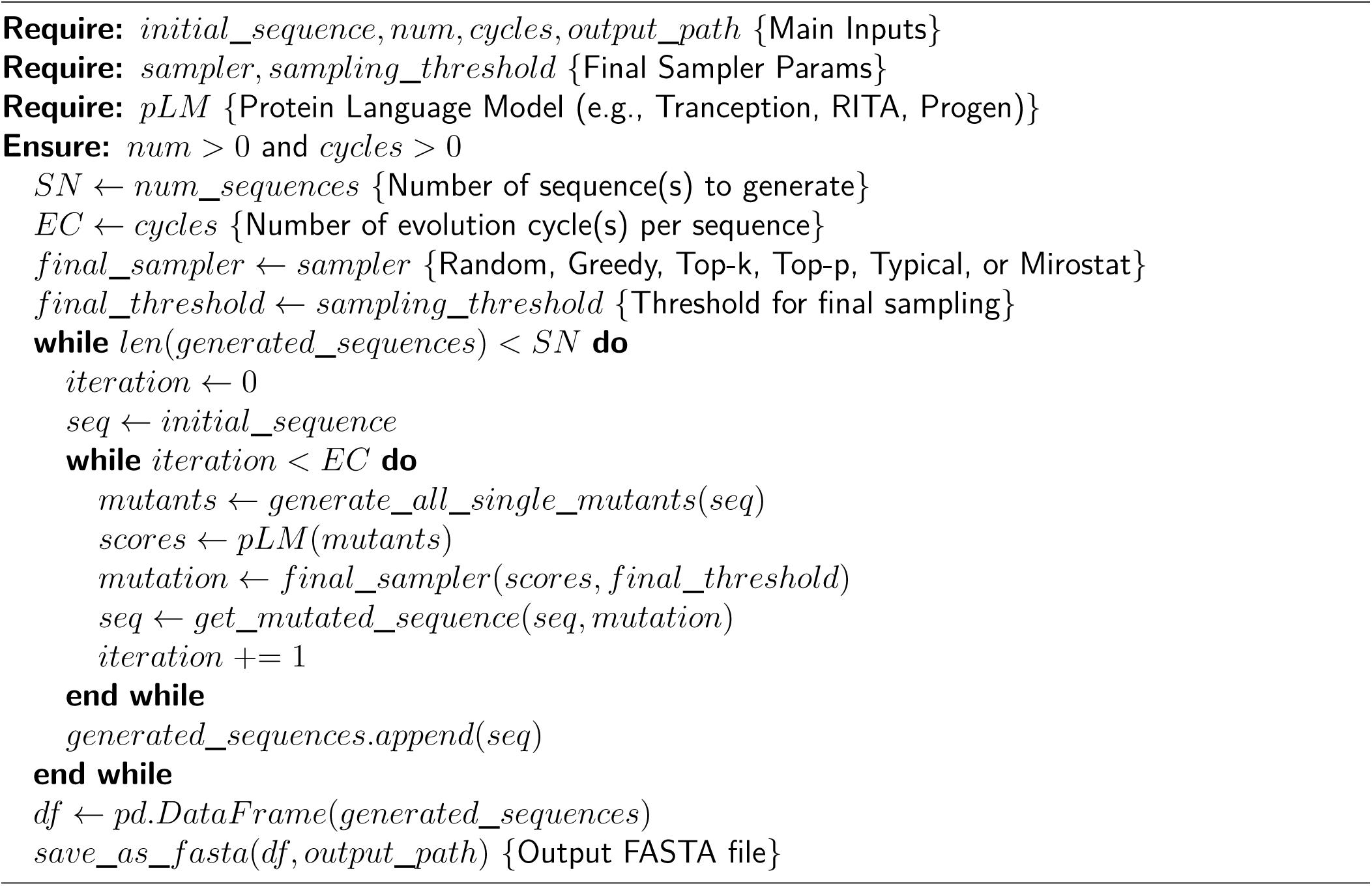

#### Algorithm S8

Autoregressive (ARS-singles and -doubles) Protein Generation Pseudocode

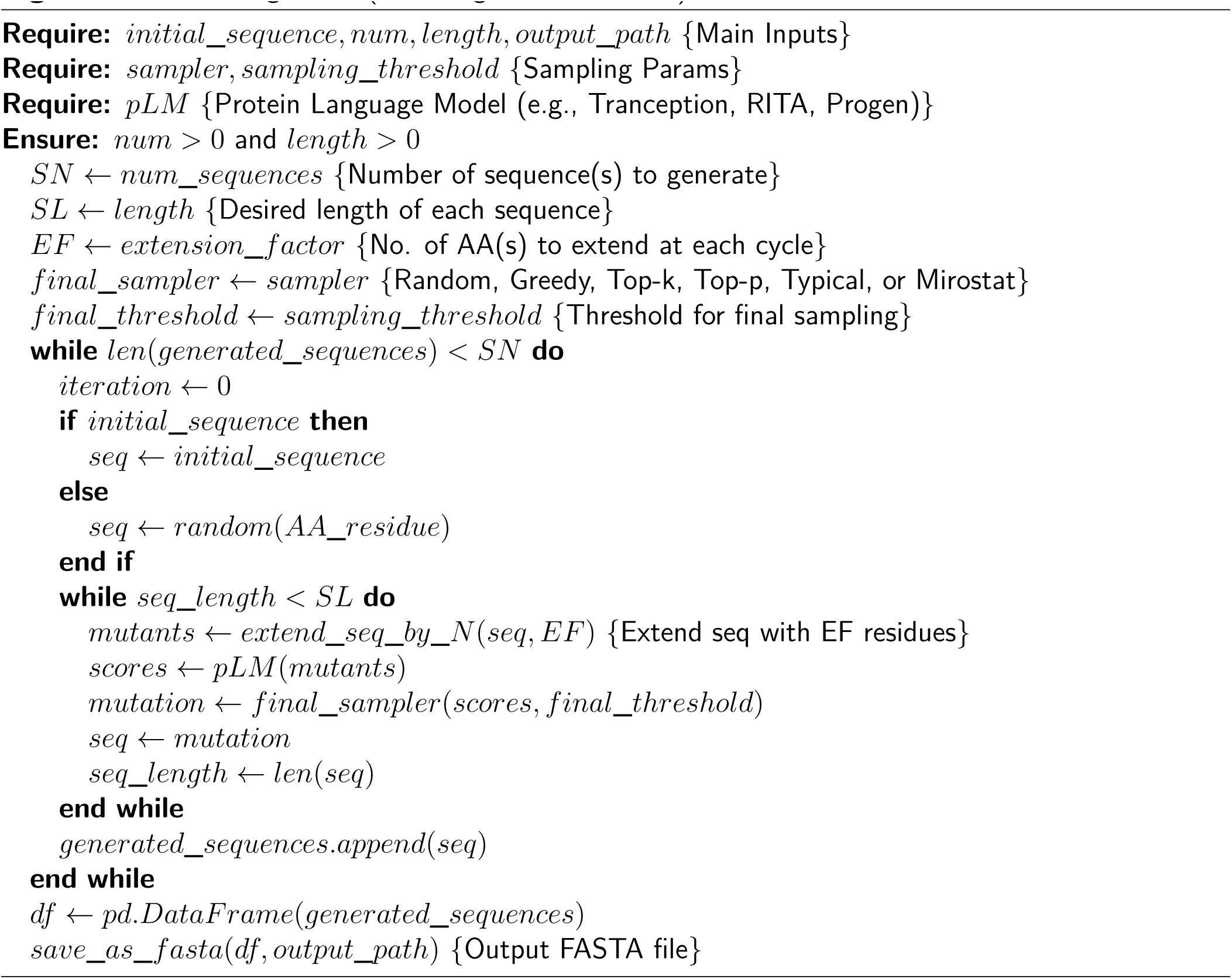

#### Algorithm S9

IRS-doubles Protein Generation Pseudocode

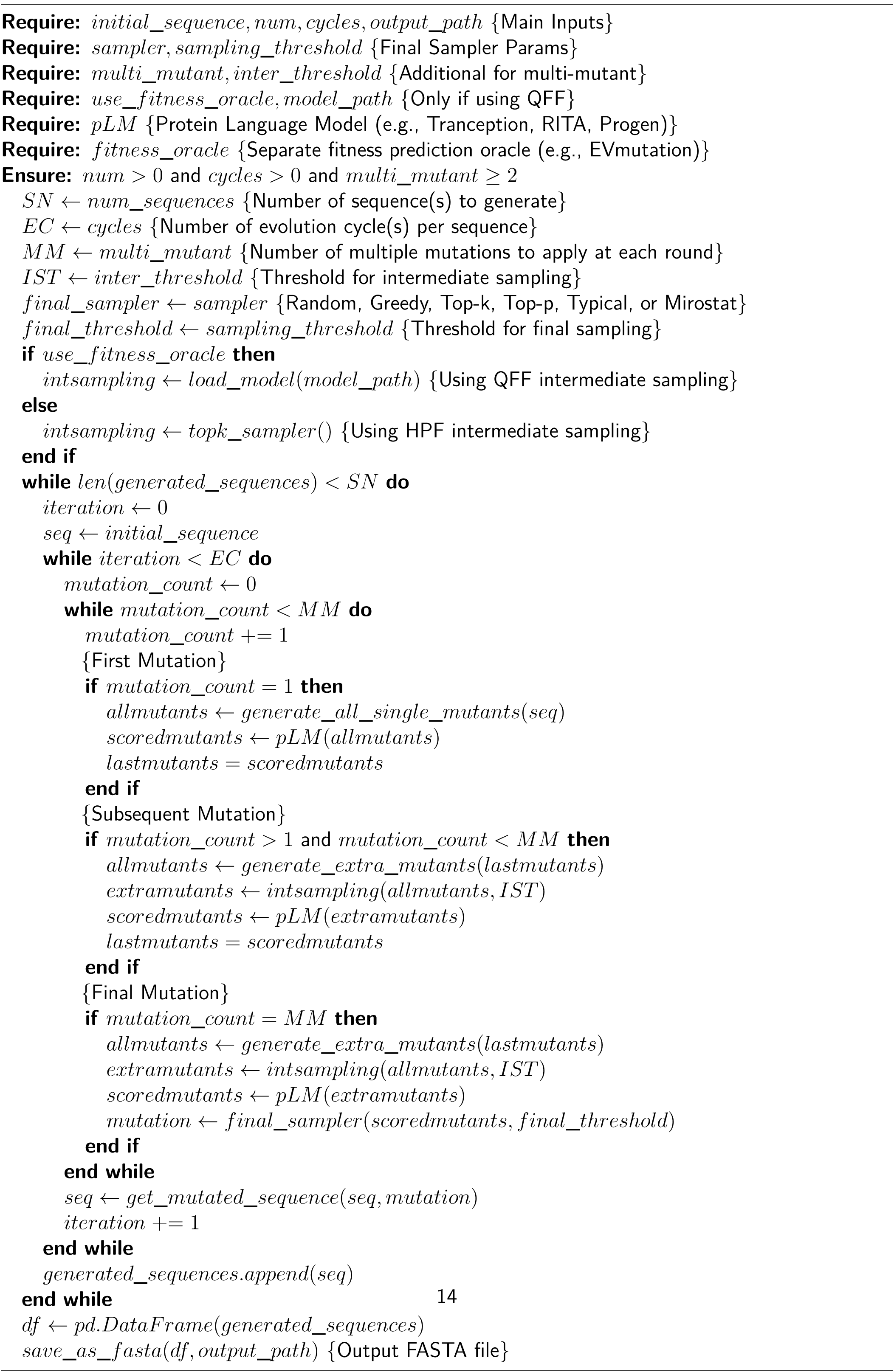

